# Evolutionarily conserved temperature dependence leads to loss of protein O-GlcNAc in mammalian hypothermia

**DOI:** 10.1101/2025.07.08.660373

**Authors:** Divya Upadhyay, Christa Kietz, Nasrin Sultana, Rishi Banerjee, Massimiliano Gaetani, Maaria Kankare, Vineta Fellman, Janne Purhonen, Jukka Kallijärvi

**Affiliations:** Folkhälsan Research Center; Haartmaninkatu 8, 00290 Helsinki, Finland; Stem Cells and Metabolism Research Program, Faculty of Medicine, University of Helsinki, P.O. Box 63, 00014 Helsinki, Finland; Chemical Proteomics, Department of Medical Biochemistry and Biophysics, Karolinska Institutet, SciLifeLab and the Swedish National Infrastructure for Biological Mass Spectrometry (BioMS), Stockholm, Sweden; Department of Biological and Environmental Science, University of Jyväskylä, P.O. Box 35, 40014, Jyväskylä, Finland; Department of Clinical Sciences, Lund, Pediatrics, Lund University, P.O. Box 117, 221 00 Lund, Sweden; Children’s Hospital, Clinicum, University of Helsinki, P.O. Box 63, 00014 Helsinki, Finland; Division of Clinical Microbiology, Department of Laboratory Medicine, Karolinska Institutet 14152 Huddinge, Sweden; Section for Neurobiology and DANDRITE, Department of Molecular Biology and Genetics, Aarhus University, Aarhus, Denmark

## Abstract

O-N-acetylglucosaminylation (O-GlcNAcylation) is a non-canonical glycosylation of intracellular proteins suggested to be involved in diverse cellular processes, yet its regulation and specific effects on protein function remain elusive. The O-GlcNAc modification of protein serine and threonine residues is cycled by only two enzymes, O-GlcNAc transferase (OGT) and O-GlcNAcase (OGA). O-GlcNAc sites lack a strict consensus sequence, suggesting that structural context rather than a defined motif governs the O-GlcNAc site selection. O-GlcNAcylation has been thought to be primarily regulated by the availability of its substrate, UDP-GlcNAc, produced by the hexosamine biosynthetic pathway (HBP). Here, we report a systemic loss of protein O-GlcNAc in *Bcs1l* mutant mice, a model of mitochondrial complex III deficiency. Although HBP precursor metabolites linked to mitochondrial function were decreased in the *Bcs1l* mutant liver, UDP-GlcNAc levels in the liver and other tissues were largely unaffected and did not correlate with the O-GlcNAc levels. Consistently, neither boosting UDP-GlcNAc with N-acetylglucosamine supplementation nor overexpression of the HBP rate-limiting enzyme GFPT1 restored O-GlcNAc levels, indicating that the loss was independent of the HBP. Instead, the body temperature of the hypothermic mutant mice showed a strong positive correlation with their O-GlcNAc level. Correcting body temperature via liver-targeted gene therapy, expression of alternative oxidase (AOX), or housing at 35°C prevented the loss of O-GlcNAc. Pharmacological inhibition of OGA restored O-GlcNAc levels in the hypothermic mice but did not affect disease progression, indicating that decreased O-GlcNAc is a physiologically normal consequence of hypothermia. We show that temperature dictates the O-GlcNAc status also across *Drosophila* species, zebrafish, and cultured mammalian cells. In cultured cells, hypothermia decreased OGT and increased OGA expression transcriptionally, indicating their ratio as a simple regulatory mechanism. Proteome solubility profiling revealed that O-GlcNAc alters the proteome’s structural and/or interactome landscape. Together, our findings identify temperature as an evolutionarily conserved master regulator of O-GlcNAcylation, which we propose protects the proteome from temperature-dependent fluctuations in both poikilothermic and homeothermic organisms.

## Introduction

O-N-acetylglucosaminylation (O-GlcNAcylation) is a dynamic co- and post-translational modification of primarily nuclear and cytosolic proteins, implicated in several cellular processes, such as gene expression, metabolism, and differentiation (Yang and Qian, 2017). O-GlcNAcylation of individual proteins has been suggested to be sensitive to cellular stress and play a cytoprotective role (Martinez et al., 2017; Zhao et al., 2024). Despite extensive literature since its discovery in the early 1980s (Holt and Hart, 1986; Torres and Hart, 1984), the physiological and biochemical mechanisms that regulate O-GlcNAc abundance remain poorly understood. Moreover, how O-GlcNAc alters protein structure and function is still unclear. Although numerous protein- or protein site-specific effects have been proposed, few nucleocytoplasmic O-GlcNAc-binding proteins or domains that would operate as readers of the modification and mediate possible downstream effects, have been identified (Toleman et al., 2018). Instead, recent chemical proteomics studies have suggested that O-GlcNAc may play a role in the regulation of protein thermostability (King et al., 2022) and protein complex solubility (Xu et al., 2025).

O-GlcNAcylation is catalyzed by O-GlcNAc transferase (OGT), which transfers N-acetylglucosamine from the activated sugar donor UDP-N-acetylglucosamine (UDP-GlcNAc), and is reversed by O-GlcNAcase (OGA) (Hardivillé and Hart, 2014; Yang and Qian, 2017; Zachara et al., 2022). UDP-GlcNAc is produced by the hexosamine biosynthetic pathway (HBP), which utilizes adenosine triphosphate (ATP), uridine triphosphate (UTP), glucose 6-phosphate, glutamine, and acetyl-CoA as precursors to produce the amino sugar. Downstream, UDP-GlcNAc is epimerized into UDP-N-acetylgalactosamine (UDP-GalNAc) for canonical glycosylation in the secretory pathway (Akella et al., 2019). As the HBP integrates nucleotide, carbohydrate, amino acid, and fatty acid metabolism, it has been proposed to function as a metabolic sensor that modulates O-GlcNAcylation primarily via UDP-GlcNAc availability. Very few studies have, however, measured the actual UDP-GlcNAc concentrations (Sunden et al., 2023). The availability of the HBP substrates, or metabolic precursors of UDP-GlcNAc, is linked directly or indirectly to mitochondrial metabolism (Akella et al., 2019; Banerjee et al., 2021). For example, *de novo* UTP biosynthesis depends on the electron transport chain (ETC)-dependent enzyme dihydroorotate dehydrogenase (DHODH). In oxidative phosphorylation (OXPHOS) deficiency, ATP production can become reliant on glycolysis, often resulting in hypoglycemia in mitochondrial diseases, while glutamine can serve as an alternative substrate for the tricarboxylic acid (TCA) cycle when glucose metabolism is disrupted (Bornstein et al., 2023). In addition, mitochondria are a major source of acetyl-CoA, depending on the metabolic state and carbon source (Banerjee et al., 2021). Together, these links suggest that O-GlcNAc homeostasis should be particularly vulnerable to mitochondrial dysfunction. It is a surprising knowledge gap that the HBP and O-GlcNAcylation have, to our knowledge, never been studied in models of primary mitochondrial diseases.

Here, we set out to investigate hexosamine biosynthesis and, as its readout, O-GlcNAcylation in a *Bcs1l^p.S78G^* knock-in mouse model of GRACILE syndrome, a neonatally lethal mitochondrial complex III (CIII) deficiency (Fellman et al., 1998; Purhonen et al., 2020; Visapää et al., 2002). We found drastically decreased global O-GlcNAc levels in these mice in both the symptomatic and non-symptomatic tissues. Rather than the HBP, we identify temperature as the master regulator of O-GlcNAc levels.

## Results

### CIII-deficient mice show global loss of protein O-GlcNAc

We quantified the levels of direct or indirect UDP-GlcNAc precursors (Fig 1A) in the blood and liver tissue of juvenile (postnatal day 30, P30) *Bcs1l^p.S78G^;mt-Cyb^p.D254N^* mice (for simplicity referred to as *Bcs1l^p.S78G^* mice from now on), a model of neonatal mitochondrial CIII deficiency. These mice carry a missense mutation in the CIII assembly factor BCS1L and a natural variant in the mtDNA-encoded CIII subunit cytochrome b, with survival of only little over one month (Purhonen et al., 2020). Blood glucose, hepatic UTP and acetyl-CoA concentration as well as glutamine/glutamate ratio were all significantly decreased in the mutant mice (Fig. 1B-E). The depletion of multiple precursors hinted at compromised hexosamine biosynthesis, possibly resulting in reduced UDP-GlcNAc production and protein glycosylation. As a readout for this, we detected protein O-GlcNAc using the RL2 monoclonal antibody and found markedly decreased levels in the mutant liver, kidney, and skeletal muscle (Fig. 1F-H). Immunohistochemical staining of liver and kidney sections showed a similar loss of O-GlcNAc, particularly abundant in the nuclei (Fig. 1I and Sup. Fig. 1A). To determine whether the loss of O-GlcNAc reflected a general defect in protein glycosylation, we probed liver, kidney and skeletal muscle lysates with wheat germ agglutinin (WGA) and concanavalin A (Con A) lectins, which recognize canonical O- and N-linked glycosylations. We found no differences in canonical glycosylation levels or patterns between WT and mutant mice (Sup. Fig. 1B-D). Together, these findings demonstrate O-GlcNAc depletion across multiple tissues in the CIII-deficient mice. We corroborated the results in adult mice carrying the *Bcs1l*-mutation in WT mtDNA background, with milder CIII deficiency and survival of 4-6 months (Purhonen et al., 2020; Rajendran et al., 2019). At P200, also these mice had a significant loss of O-GlcNAc in the liver, kidney and especially in the skeletal muscle (Sup. Fig. 1E-G).

**Figure 1.**
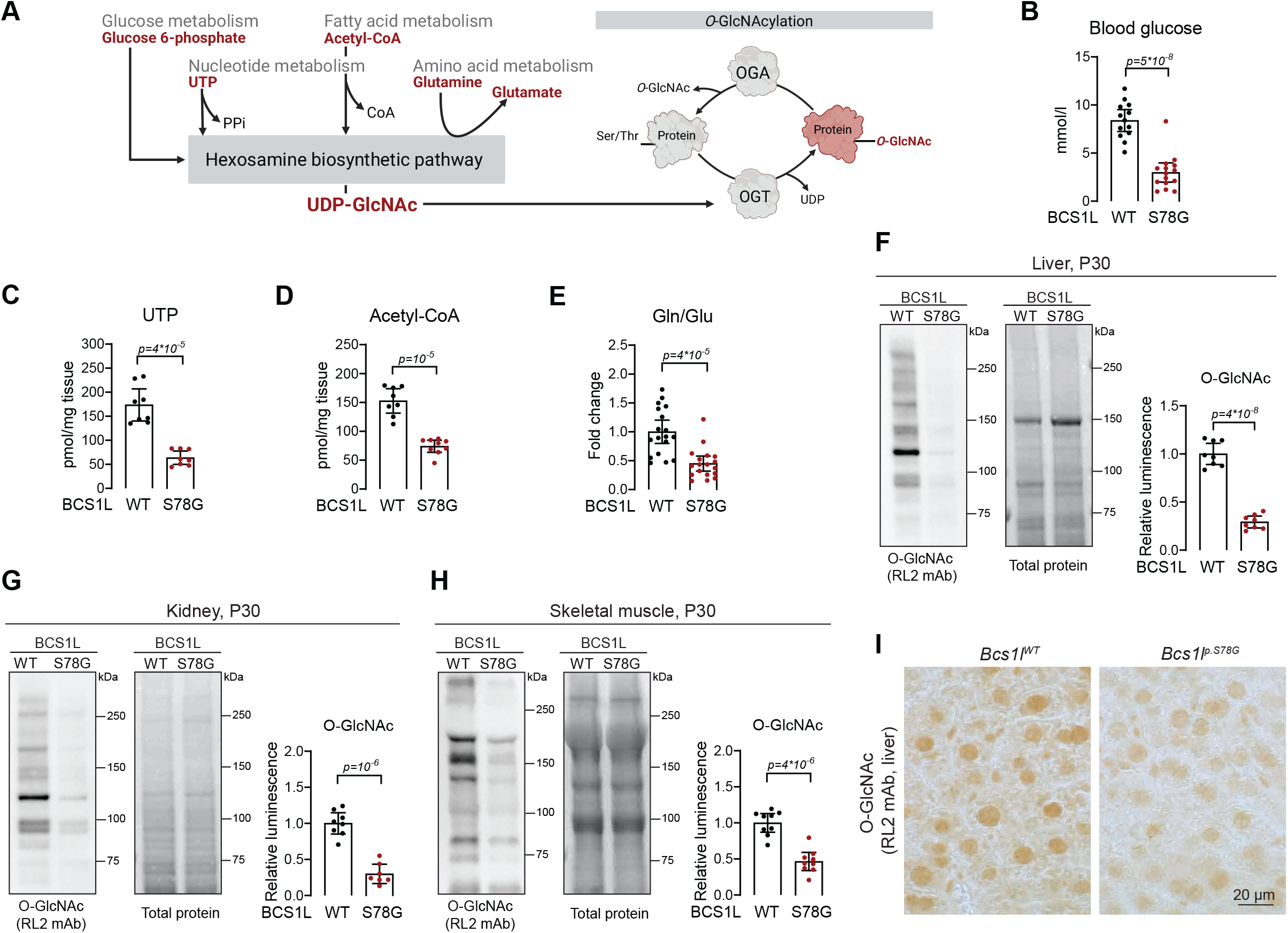
Global loss of protein O-GlcNAc in CIII-deficient *Bcs1l^p.S78G^* mice. A) Schematic illustration of the hexosamine biosynthetic pathway and O-GlcNAcylation. B) Blood glucose concentration in WT and *Bcs1l^p.S78G^* mice at P30, n = 13-15/genotype. C) Liver UTP concentration, n = 8/genotype. D) Liver acetyl-CoA concentration, n = 8-9/genotype. E) Ratio of glutamine to glutamate in liver tissues, n = 18/genotype. F-H) Representative western blots and quantifications of O-GlcNAc levels in the liver (F), kidney (G), and skeletal muscle (H) lysates, n = 7-9/genotype. I) Representative O-GlcNAc immunohistochemistry image of liver sections. Statistics: Welch’s two-sided t-test. *p* values of the group comparisons are indicated above the panel. The error bars represent 95% CI of mean. All data points derive from independent mice. All mice were on the *mt-Cyb^p.D254N^*background.

### Neither compromised hexosamine biosynthesis nor UDP accumulation explains the loss of O-GlcNAc in CIII deficiency

To assess if decreased UDP-GlcNAc availability contributed to the loss of O-GlcNAc, we first performed LC-MS quantification of UDP-N-acetylhexosamines (UDP-GlcNAc + UDP-GalNAc) in liver samples. These metabolites were unchanged between WT and mutant mice (Sup. Fig. 1H). Using our previously developed enzymatic assay to specifically quantify UDP-GlcNAc (Sunden et al., 2023; Upadhyay et al., 2024), we found no consistent decrease in UDP-GlcNAc across the liver, kidney, and skeletal muscle of the *Bcs1l^p.S78G^* mice (Fig. 2A). The liver showed a slight statistically significant decrease in UDP-GlcNAc and a weak correlation with protein O-GlcNAc (R²=0.17) but there was no correlation in the kidney and skeletal muscle (Fig. 2B). To test whether increasing UDP-GlcNAc availability through the salvage pathway could restore O-GlcNAc levels, we supplemented mouse chow with 1% (w/w) N-acetylglucosamine, together with 0.5% triacetyluridine, which serves as a precursor for UTP to bypass the potentially compromised pyrimidine nucleotide biosynthesis in the mutant mice. The glucosamine-uridine supplement from weaning to P29 left UDP-GlcNAc and O-GlcNAc levels unchanged (Fig. 2C, D). Similarly, overexpressing GFPT1, the rate-limiting enzyme of HBP, in the liver using recombinant adeno-associated viral vectors (rAAV) had no effect on hepatic UDP-GlcNAc or O-GlcNAc levels (Fig. 2E, 2F).

**Figure 2.**
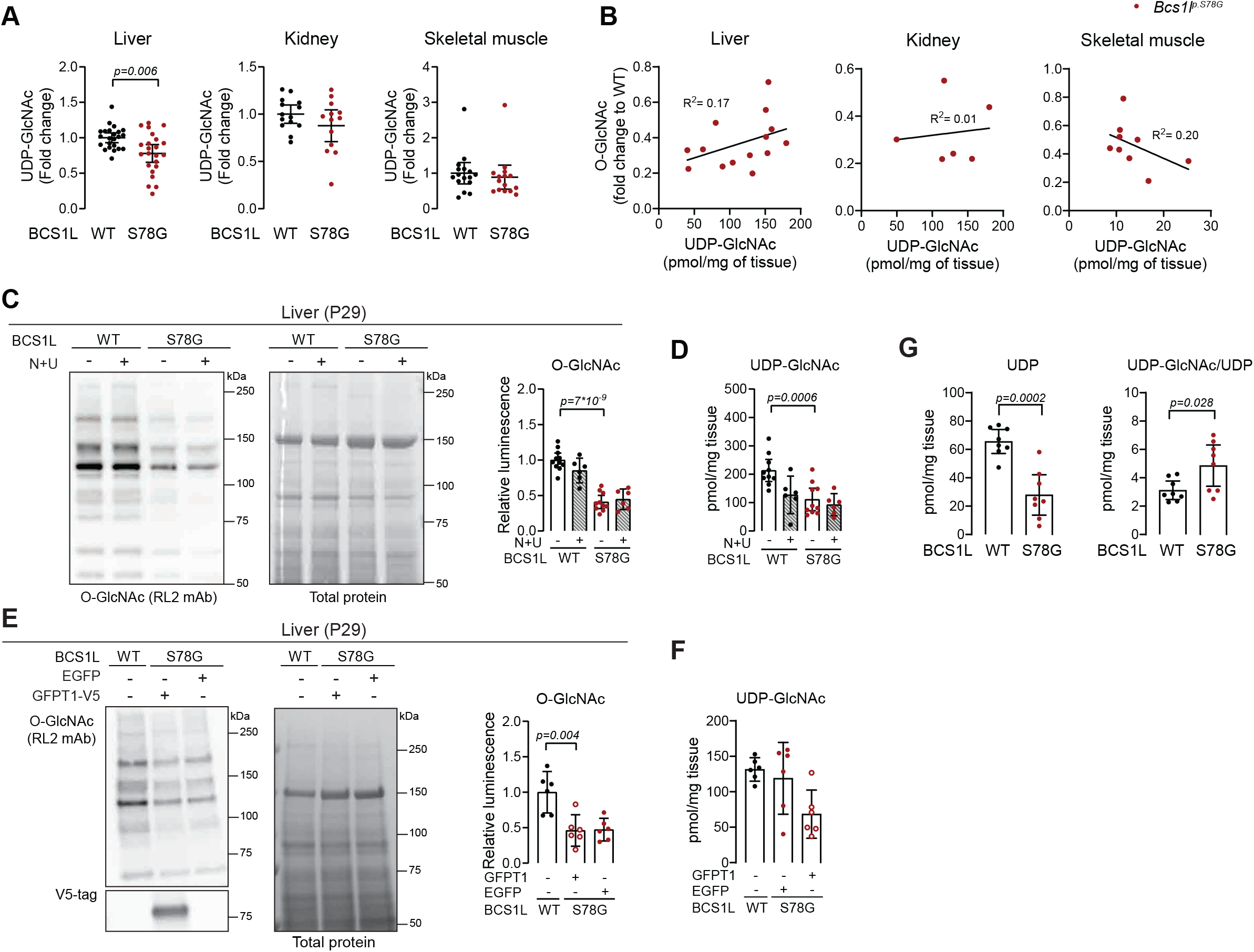
O-GlcNAc loss in *Bcs1l* mutant mice is not driven by limited UDP-GlcNAc availability or UDP-mediated inhibition of OGT. A) Concentration of UDP-GlcNAc in the liver, kidney, and skeletal muscle of WT and *Bcs1l^p.S78G^* mice, n = 13-23/genotype. B) Correlation between O-GlcNAc levels (western blot quantifications) and UDP-GlcNAc concentration in the liver, kidney and skeletal muscle, n = 6-14/genotype. C) Representative western blot and quantification of O-GlcNAc levels in the liver of WT and *Bcs1l^p.S78G^* mice fed standard chow or chow supplemented with 1% (w/w) N-acetylglucosamine and 0.5% triacyl uridine, n = 6-11/genotype. D) Concentration of UDP-GlcNAc in the liver of WT and *Bcs1l^p.S78G^* mice fed standard chow or chow supplemented with 1% (w/w) N-acetylglucosamine and 0.5% triacyluridine, n = 6-11/genotype. E) Representative western blot and quantification of O-GlcNAc levels in liver of WT mice or of *Bcs1l^p.S78G^* mice injected with *EGFP*- or *Gfpt1-V5* rAAVs, n = 6/genotype. F) Concentration of UDP-GlcNAc in the liver of WT mice or of *EGFP*- or *Gfpt1-V5-*rAAV-injected *Bcs1l^p.S78G^* mice, n = 6/genotype. G) Concentration of UDP and UDP-GlcNAc/UDP ratio in liver tissue, n = 8/genotype. Statistics: Welch’s two-sided t-test (A, G) or one-way ANOVA followed by the selected pairwise comparisons with Welch’s statistics (C-F). *p* values of the group comparisons are indicated above the panel. The error bars represent 95% CI of mean. All data points derive from independent mice. All mice were on *mt-Cyb^p.D254N^*background.

OXPHOS deficiency can compromise biosynthesis and phosphorylation of ribonucleosides (Purhonen et al. 2023a), potentially including accumulation of UDP, which is a strong end-product inhibitor of OGT (Sunden et al., 2023). We quantified UDP and UTP in the mutant liver and found no UDP accumulation but instead depletion by about 60% (Fig. 2G), leading to increased UDP-GlcNAc/UDP ratio (Fig. 2G). Together, these findings indicate that the loss of protein O-GlcNAc in the CIII-deficient mice was not driven by decreased UDP-GlcNAc availability or UDP-mediated inhibition of OGT.

### Temperature is a conserved regulator of global O-GlcNAc level across animal species

Having excluded UDP-GlcNAc availability as the cause of O-GlcNAc loss in CIII deficiency, we sought alternative mechanisms. We recently discovered that *Bcs1l^p.^*^S78G^ mice develop hypothermia due to insufficient basal and adaptive thermogenesis (preprint Banerjee et al., 2024). Intriguingly, a *Drosophila* study published in 2014 showed positive correlation between O-GlcNAc level and ambient temperature during embryonic development of this poikilothermic invertebrate species, although the mechanism and purpose of the phenomenon remained unknown (Radermacher et al., 2014). We revisited and extended this observation to mammalian systems and analyzed O-GlcNAc in mouse hepatocyte-derived cell lines AML12 and Hepa1-6, and in human dermal fibroblasts cultured at temperatures of 30 °C, 37 °C and 41 °C. Indeed, also in cells from these homeothermic mammals, temperature strongly determined the O-GlcNAc abundance (Fig. 3A). In the hepatocyte-derived cells, O-GlcNAc levels decreased gradually with colder temperature in a time-dependent manner and remained low upon extended culture of five days at 30 °C (Fig. 3B-D, Sup. Fig. 2A, B). In contrast to O-GlcNAc levels, temperature shifts did not change overall O- and N-linked glycan levels (Sup. Fig 2C). Furthermore, UDP-GlcNAc levels remained largely stable across temperatures, except following the temperature shift to 41 °C in Hepa1-6 cells and in human fibroblasts (Fig 3E). To assess the evolutionary universality of these findings, we analyzed O-GlcNAc in the poikilothermic vertebrate model zebrafish and in adult fruit flies. In zebrafish muscle tissue, O-GlcNAc levels increased with aquarium water temperature (between 16 °C and 32 °C) (Fig. 3F). Similarly, in adult fruit flies maintained at 18 °C, 25 °C or 32 °C for 16 hours, O-GlcNAc levels increased progressively as the temperature increased (Fig. 3G), while UDP-GlcNAc concentrations remained unchanged (Fig. 3H). To extend this observation to a cold-adapted insect species, we analyzed *Drosophila montana* maintained at temperatures spanning −6 °C to +19 °C (Hoikkala and Poikela, 2022). O-GlcNAc was abundant at 19 °C but almost undetectable at 10 °C and below (Fig. 3I). Together, these data identify temperature as a conserved determinant of global O-GlcNAc abundance across poikilothermic and homeothermic species.

**Figure 3.**
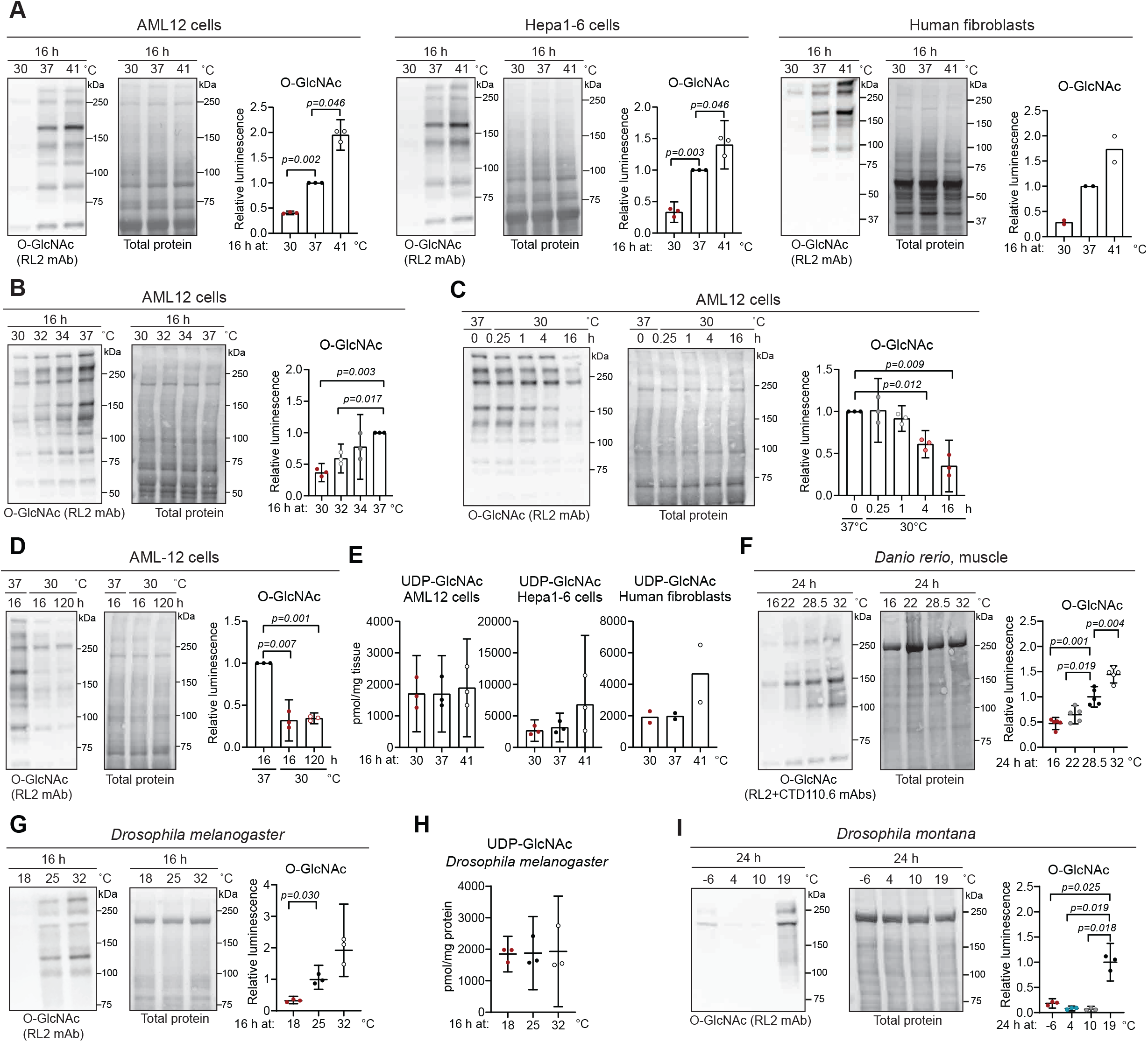
O-GlcNAc levels are temperature-dependent in mammalian cells, in the fruit fly, and in zebrafish. A) Representative western blot and quantification of O-GlcNAc levels in AML12 cells, Hepa1-6 cells, and human fibroblasts incubated 16 h at indicated temperatures. B) Representative western blot and quantification of O-GlcNAc levels in AML12 cells incubated 16 h at indicated temperatures. C) Representative western blot and quantification of O-GlcNAc levels in AML12 cells incubated for the indicated time points at 30 °C. D) Representative western blot and quantification of O-GlcNAc levels in AML12 cells maintained for the indicated time points at 30 °C. E) Concentration of UDP-GlcNAc in AML12 cells, Hepa1-6 cells, and human fibroblasts, kept 16 h at indicated temperatures. F) Representative western blot and quantification of O-GlcNAc levels in muscle tissue from zebrafish kept 24 h at indicated temperatures, n = 5. G) Representative western blot and quantification of O-GlcNAc levels in *Drosophila melanogaster* whole-body lysates from the *w^1118^* fly line kept 16 h at indicated temperatures, n = 3. H) Concentration of UDP-GlcNAc in *Drosophila melanogaster*, n = 3, kept 16 h at indicated temperatures. I) Representative western blot and quantification of O-GlcNAc levels in *Drosophila montana* whole-body lysates from WT flies kept 24 h at indicated temperatures, n = 3. Statistics: One-sample t-test (A-D) and one-way ANOVA followed by the selected pairwise comparisons with Welch’s statistics (F, G, I). *p* values of the group comparisons are indicated above the panel. Error bars represent 95% CI of mean. Data in A–E derive from three independent experiments, except for the fibroblast experiments in A and E (n = 2). In panel F, each data point derives from distinct animals. In G-I each data point represents 3-5 pooled *Drosophila* adult male flies.

### Hypothermia causes the loss of O-GlcNAc in *Bcs1l^p.S78G^* mice

In the hypothermic *Bcs1l^p.S78G^* mice, O-GlcNAc levels correlated positively with their temperature, as assessed by non-invasive infrared thermometry (Fig. 4A). To obtain direct proof for the temperature dependency of O-GlcNAc in mice, we utilized samples from *Bcs1l^p.^*^S78G^ mice, in which we restored normothermia via various interventions. First, we restored CIII function in the juvenile liver with rAAVs expressing wild-type *Bcs1l* under a hepatocyte-specific promoter, which restores body temperature without correcting CIII deficiency in other tissues (Fig. 4B) (preprint Banerjee et al., 2024). O-GlcNAc levels were normalized to WT level not only in the liver but also in the untargeted kidney (Fig. 4C). Next, we crossed the *Bcs1l^p.S78G^* mice with transgenic mice expressing a non-mammalian mitochondrial respiratory enzyme alternative oxidase (AOX), which directly and indirectly increases heat generation (El-Khoury et al., 2022; Purhonen et al., 2023a; Rajendran et al., 2019). Transgenic AOX expression warms the juvenile *Bcs1l^p.S78G^* mice up to near-wild-type core temperatures (preprint Banerjee et al., 2024). Indeed, AOX attenuated the loss of O-GlcNAc in the liver and completely prevented it in the kidney and skeletal muscle (Fig. 4D). Finally, short-term housing of the mutant mice at 35°C from P25 until sample collection at P28 forced the body temperature to near-normal (preprint Banerjee et al., 2024) and prevented the loss of O-GlcNAc in the liver and skeletal muscle (Fig. 4E, F). O-GlcNAc levels in the WT and mutant mice correlated with temperature in this experiment as well (Fig. 4G). Together, these data confirm hypothermia as the cause of the systemic loss of O-GlcNAc in the CIII-deficient mice.

**Figure 4.**
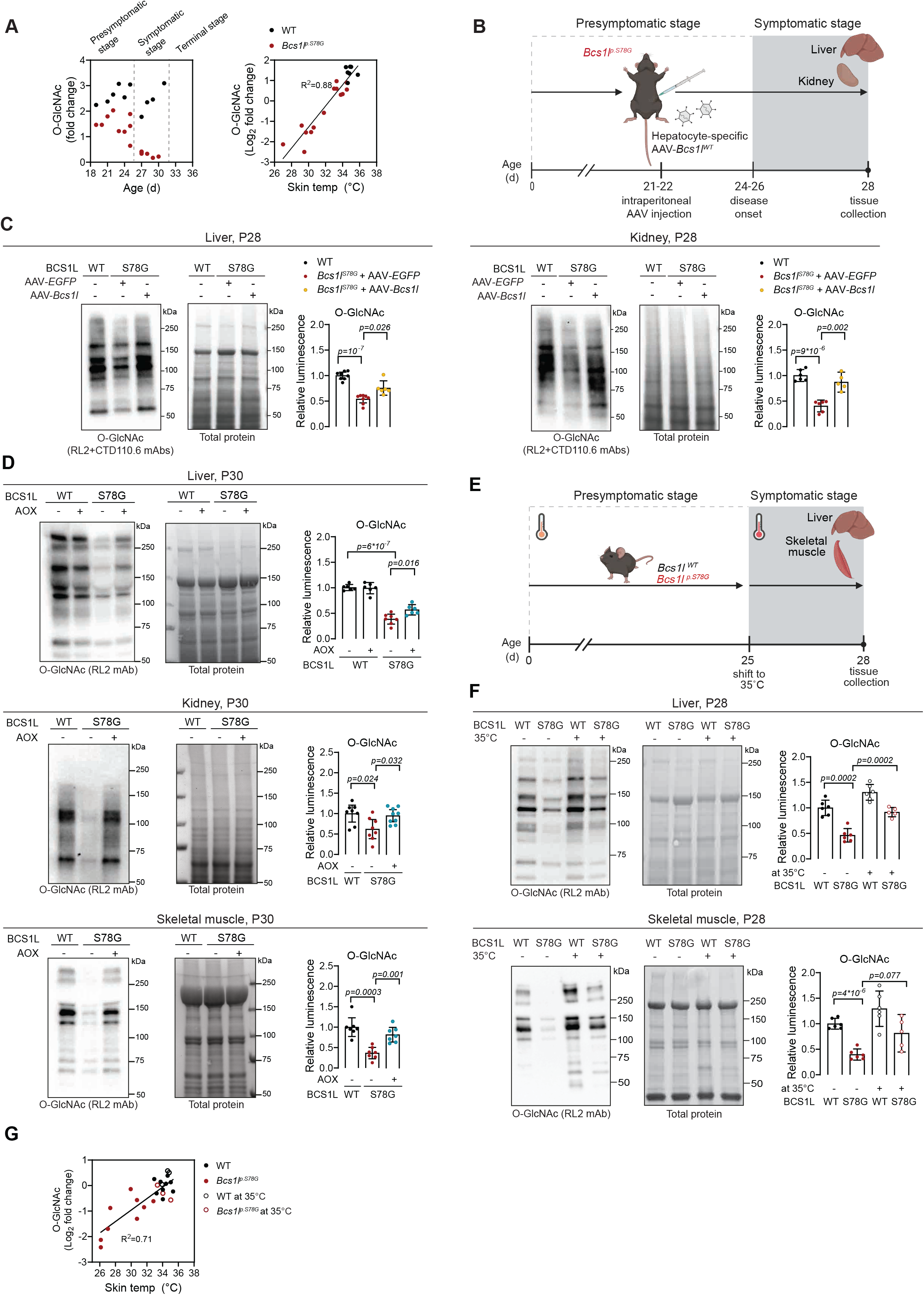
Restoration of body temperature rescues O-GlcNAc levels in *Bcs1l^p.S78G^* mice. A) Hepatic O-GlcNAc levels (western blot quantifications) plotted against mouse age and correlation between the O-GlcNAc levels and skin temperature. B) Schematic of the experimental design for rAAV-mediated liver-targeted *Bcs1l* rescue. C) Representative western blot and quantification of O-GlcNAc levels in liver and kidney tissues of WT, and *Bcs1l^p.S78G^* mice injected with rAAVs delivering hepatocyte-specific expression of *EGFP* (control) or wild-type *Bcs1l*, n = 5-6/genotype. D) Representative western blot and quantification of O-GlcNAc levels in liver tissue of WT*, AOX, Bcs1l^p.S78G^*, and *Bcs1l^p.S78G^;AOX* mice and in kidney and skeletal muscle tissues of WT*, Bcs1l^p.S78G^*and *Bcs1l^p.S78G^;AOX* mice, n = 6-9/genotype. E) Schematic of the experimental design for forced normothermia (P25–P28). F) Representative western blot and quantification of O-GlcNAc levels in liver and skeletal muscle tissues of mice kept three days at 35 °C, n = 5-6/condition. G) Correlation between hepatic O-GlcNAc levels (western blot quantification) and skin temperature. Statistics: One-way ANOVA followed by the selected pairwise comparisons with Welch’s statistics. *p* values of the group comparisons are indicated above the panel. Error bars represent 95% CI of mean. All data points derive from independent mice. All mice were on *mt-Cyb^p.D254N^* background.

### OGT and OGA transcription set O-GlcNAc levels to match temperature

According to literature, OGT and OGA activities are under tight regulation to maintain appropriate O-GlcNAc levels (Issad et al., 2022; Zhang et al., 2014). To assess if enzyme activities directly underlay the loss of O-GlcNAc in cold, we measured the activities of purified OGT and OGA as a function of temperature. As expected, the activities of both enzymes correlated with reaction temperature (Sup. Fig. 3A, B). Temperature rise from 30 to 40 °C increased OGA activity relatively more than the activity of OGT. Curiously, bacterially expressed purified OGA remained inactive at 30 °C without prior pre-incubation at 37 °C, suggesting its inefficient folding at low temperatures (Sup. Fig. 3A, B). Nevertheless, as these observations pointed to a higher rather than lower OGT/OGA activity ratio in lower temperatures, they cannot explain the positive temperature dependency of O-GlcNAc levels *in vivo*. We next quantified endogenous OGA and OGT proteins in AML12 (Fig. 5A) and Hepa1-6 cells (Sup. Fig. 3C) at different temperatures. In contrast to most perturbations of O-GlcNAcylation, which typically lead to a robust compensatory shift in OGT/OGA balance (Sunden et al., 2023), low temperature instead increased OGA and decreased OGT protein expression, indicating lack of compensation but rather active downregulation of O-GlcNAc levels. Thus, temperature affects O-GlcNAcylation via the OGT/OGA ratio (Fig. 5A). In AML12 cells stably expressing tagged OGT and OGA from chicken β-actin promoter, cold exposure increased endogenous OGA and decreased endogenous OGT levels but did not affect the levels of the tagged proteins, suggesting transcriptional regulation (Fig. 5B).

**Figure 5.**
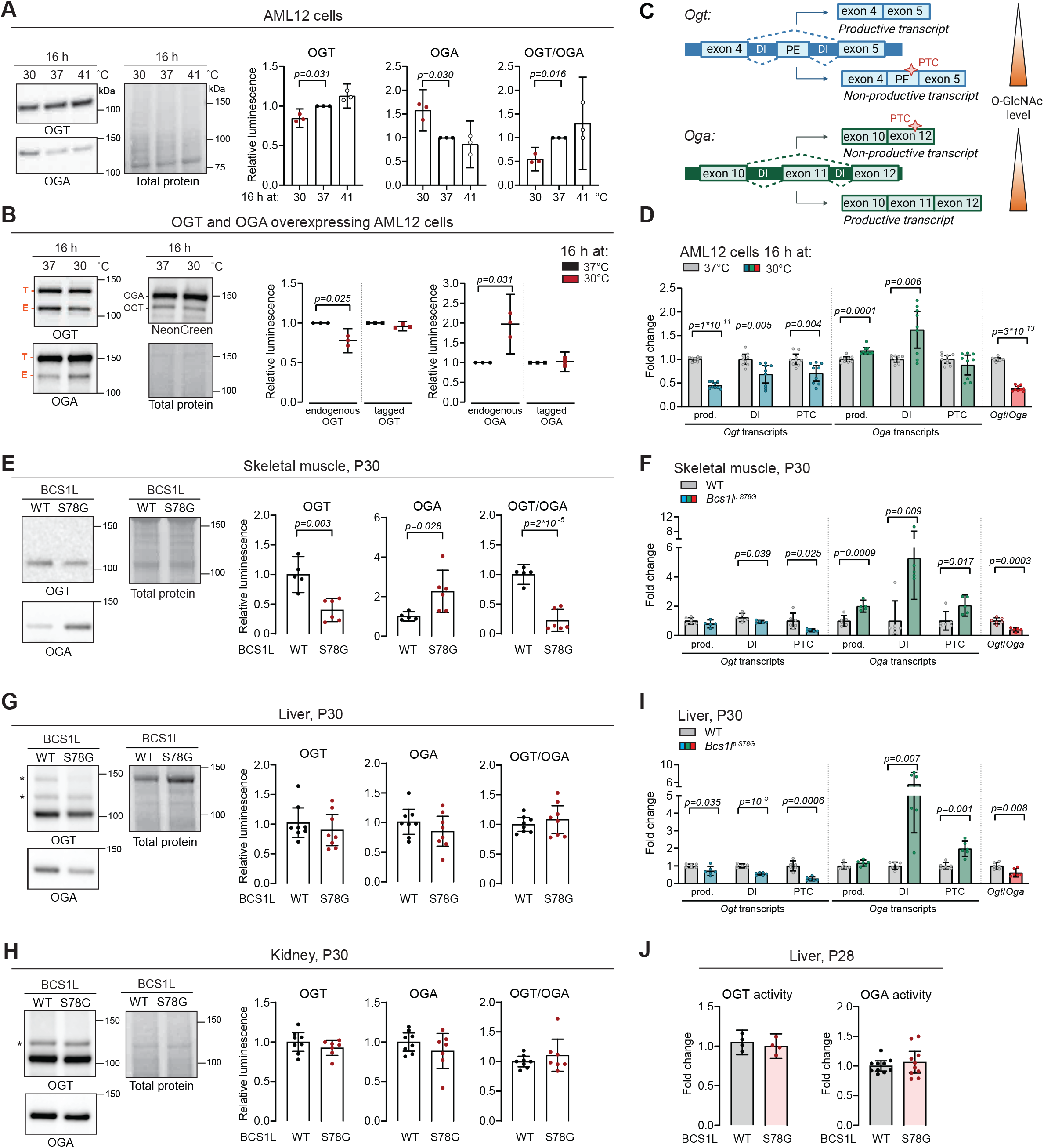
Effect of temperature on OGT and OGA expression. A) Representative western blot and quantification of OGT and OGA protein levels, and their ratio, in AML12 cells incubated 16 h at indicated temperatures, n = 3 experiments B) Representative western blot and quantification of endogenous (E) and NeonGreen-tagged (T) OGT and OGA in AML12 cells incubated 16 h at indicated temperatures, n = 3 experiments C) Schematic illustration of detained intron (DI)-splicing mechanism regulating expression of *Ogt* and *Oga* transcripts according to O-GlcNAc levels. D, F, I) mRNA expression of the productive, DI-containing and premature termination codon (PTC)-containing transcripts of *Ogt* and *Oga* as well as the ratio of the productive transcript of *Ogt* and *Oga* in AML12 cells kept 16 h at the indicated temperatures (D) and in skeletal muscle (F) or liver (I) of WT and *Bcs1l^p.S78G^* mice, n = 5-6/genotype. E, G, H) Representative western blot and quantification of OGT and OGA protein levels, and their ratio, in skeletal muscle (E), liver (G), and kidney (H) tissue of WT and *Bcs1l^p.S78G^* mice, n = 5-6/genotype. J) OGT and OGA activity in P28 liver tissue, n = 4 or 10/genotype, respectively. Statistics: One-sample t-test (A, B), one-way ANOVA followed by the selected pairwise comparisons with Welch’s statistics (D, F, I). Welch’s two-sided t-test (E, G, H). *p* values of the group comparisons are indicated above the panel. The error bars represent 95% CI of mean. Data in A, B, and D derive from three independent experiments. Each data point in E-J derives from independent mice. All mice were on *mt-Cyb^p.D254N^* background. *, isoforms of OGT or unspecific bands.

Perturbations in O-GlcNAcylation have been shown to trigger a detained intron (DI)-splicing feedback mechanism that adjusts OGT and OGA expression in cultured cells (Fig. 5C) (Tan et al., 2020). To learn whether hypothermia engages this mode of regulation, we quantified the productive, DI-containing, and premature termination codon (PTC)-containing *Ogt* and *Oga* transcripts in hypothermic cells. We found that hypothermic cells expressed reduced levels of all *Ogt* transcripts, indicating that hypothermia does not activate the DI-splicing feedback despite reduced O-GlcNAc levels (Fig. 5D). In contrast, hypothermic cells increased both the productive and DI-containing *Oga* transcripts, consistent with active upregulation of *Oga* rather than a compensatory response to restore O-GlcNAc levels. Consistent with the protein data, hypothermic cells showed a decreased ratio of productive *Ogt* to *Oga* transcripts, favoring *Oga* expression (Fig. 5D). In the hypothermic mutant skeletal muscle, OGT protein was decreased and OGA increased (Fig. 5E). All *Ogt* transcripts were reduced, whereas productive and DI-containing *Oga* transcripts increased, resulting in a reduced ratio of productive *Ogt* to *Oga* transcripts, resembling the response seen in the cells cultured in hypothermic conditions (Fig. 5F).

In contrast to the skeletal muscle, neither OGT nor OGA protein levels changed significantly in the mutant liver or kidney despite the marked reduction in O-GlcNAc (Fig. 5G, H). Hepatic transcript analysis nevertheless showed reduced expression of all *Ogt* transcript isoforms, increased DI- and PTC-containing *Oga* transcripts, whereas the productive *Oga* transcript remained unchanged, resulting in a reduced ratio of productive *Ogt*/*Oga* transcripts (Fig. 5I). Thus, many transcriptional features of the hypothermic response were preserved in the liver, but these changes were not accompanied by altered OGT or OGA protein abundance, possibly due to compensation for the slightly decreased UDP-GlcNAc levels (Sunden et al., 2023). Consistent with the unchanged protein levels, immunoprecipitated OGT from mutant livers showed normal activity toward a peptide substrate, and OGA activity in the mutant liver tissue lysates was also unchanged (Fig. 5J). Similar tissue-specific patterns of OGT and OGA were observed in adult (P200) mice with the WT mtDNA background (Sup. Fig. 3D-F). In contrast, skeletal muscle OGT/OGA ratio correlated with core temperature in these mice already at P33 (Sup. Fig. 3G).

### High O-GlcNAc levels are dispensable in hypothermia

To test if the hypothermic O-GlcNAc loss is an active process, we inhibited OGA-mediated O-GlcNAc removal with thiamet G (TMG) in the AML12 cells cultured at 30 °C. As a second strategy, we employed a viable, hypomorphic mutant *Oga* allele in fruit flies (*Drosophila melanogaster*) and kept the mutant and wildtype flies at 6 °C or 18 °C for 16 h. These manipulations prevented the loss of O-GlcNAc, demonstrating that the hypothermia-induced O-GlcNAc loss is mediated by active OGA-dependent removal rather than reduced O-GlcNAc transfer alone or protein turnover (Fig. 6A, B, Sup. Fig. 4A).

**Figure 6.**
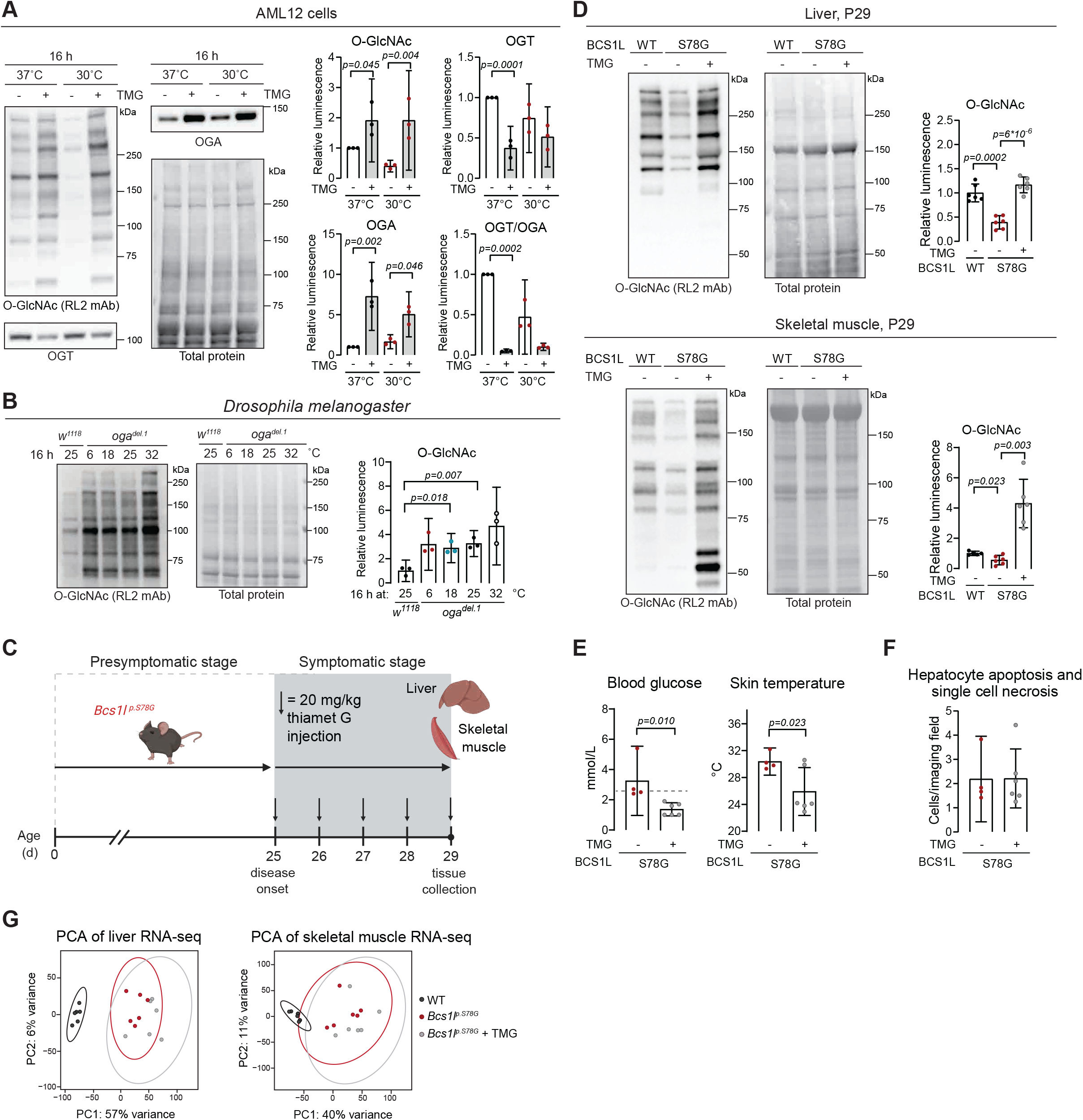
Enhancing O-GlcNAc levels in the hypothermic mice lacks therapeutic effect. A) Representative western blot and quantification of O-GlcNAc, OGT and OGA protein levels in AML12 cells ± 0.5 µM Thiamet G (TMG) incubated 16 h at 30 °C or 37 °C, n = 3 experiments. B) Representative western blot and quantification of O-GlcNAc levels in *w^1118^*and *oga^del.1^* fly lines kept 16 h at indicated temperatures, n = 3. C) Schematic of experimental design for TMG treatment. D) Representative western blot and quantification of O-GlcNAc levels in liver and skeletal muscle tissue of WT, *Bcs1l^p.S78G^;mt-Cyb^p.D254N^,* and TMG-injected *Bcs1l^p.S78G^* mice, n = 6/condition. E) Blood glucose concentration and skin temperature of *Bcs1l^p.S78G^* and TMG-injected *Bcs1l^p.S78G^* mice, n = 4-6/genotype. F) Number of apoptotic and necrotic cells in WT, *Bcs1l^p.S78G^* and TMG-injected *Bcs1l^p.S78G^* mice. G) Principal component analysis of liver and skeletal muscle transcriptomics data set from WT, *Bcs1l^p.S78G^,* and TMG-injected *Bcs1l^p.S78G^* mice. Statistics: Two-way ANOVA followed by Šídák’s multiple comparisons test for selected pairwise comparisons (A), one-way ANOVA followed by the selected pairwise comparisons with Welch’s statistics (B, D), Mann-Whitney U test (E, blood glucose), and Welch’s two-sided t-test (E, skin temperature). *p* values of the group comparisons are indicated above the panel. Error bars represent 95% CI of mean. In A, the data are derived from three independent experiments. In B, each data point consists of 3-5 pooled *Drosophila* adult male flies. Each data point in D-G derives from independent mice. All mice were on *mt-Cyb^p.D254N^* background.

To gain insight into the consequences of the hypothermic O-GlcNAc loss in cells from a homeothermic mammal, we performed transcriptomics and proteomics from the cold- and TMG-treated AML12 cells. Strikingly, restoring O-GlcNAc during hypothermia with TMG had minimal effect on the transcriptome and proteome (Sup. Fig. 4B, 4C). Next, to determine whether hypothermia-induced O-GlcNAc loss contributed to disease progression in the *Bcs1l^p.S78G^* mice, we inhibited OGA *in vivo* with daily TMG i.p. injections (Fig. 6C). After four days, hepatic O-GlcNAc levels were at WT levels while the skeletal muscle of the TMG-treated mutant mice had even two-fold higher O-GlcNAc levels compared to that of WTs (Fig. 6D). Unexpectedly, TMG treatment tended to further lower the blood glucose and temperature of the mutant mice (Fig. 6E), indicating that O-GlcNAc level was irrelevant to the disease progression. The liver pathology was not improved either, as the increased number of apoptotic or necrotic cells remained unchanged following TMG treatment (Figure 6F). Again, liver and skeletal muscle transcriptomes from untreated and TMG treated *Bcs1l^p.S78G^*mice showed minimal differences (Fig. 6G). Taken together, the cell and mouse data show that O-GlcNAc abundance decreases at lower temperatures, consistent with a temperature-dependent regulation that maintains O-GlcNAc levels within an appropriate physiological range.

### Hypothermic loss of O-GlcNAc leads to proteome solubility changes

Because restoring O-GlcNAc levels during hypothermia produced no mechanistic clues about the purpose of its temperature dependence (Sup. Fig. 4B), we next inhibited OGT in AML12 cells maintained at 37 °C. As OGT is essential for cell survival (Slawson et al., 2005) we selected a concentration of the OGT inhibitor OSMI-1 that reduced global O-GlcNAc levels by approximately 40% (Sup. Fig. 4E). Despite this modest reduction, 16 hours of OGT inhibition had a robust effect on the transcriptome (Fig. 7A). Gene set enrichment analysis (GSEA) identified the Hallmark Unfolded Protein Response as the second most significantly enriched gene sets in OSMI-1-treated cells (Fig. 7A), suggesting a role for O-GlcNAcylation in proteostasis. Subsequently, we hypothesized that, rather than being a metabolic regulator, O-GlcNAcylation confers a proteome-wide structural adaptation to temperature. To explore this, we performed a Proteome Integral Solubility Alteration (PISA) assay to assess whether hypothermic loss of O-GlcNAc alters protein solubility after a challenge with the PISA temperature gradient. TMG-treated cells grown at 30 °C (with high O-GlcNAc level) and cells grown at 37 °C served as references. Although a large fraction of proteins from hypothermic cells showed enhanced resistance to aggregation and/or precipitation compared with proteins from cells cultured at 37 °C, a subset of proteins showed an opposite trend (Fig. 7B). In this subset, TMG produced resistance to thermal aggregation in 25% of proteins (Fig. 7B, C, Sup. Table 1). In contrast, only 6% of proteins that were more soluble in hypothermic cells compared with 37 °C also remained more soluble compared to TMG-treated cells (Fig. 7B, Sup. Table 2). Together, these findings suggest that reduced O-GlcNAc levels during hypothermia are associated with selective changes in protein solubility rather than proteome-wide expression changes.

**Figure 7.**
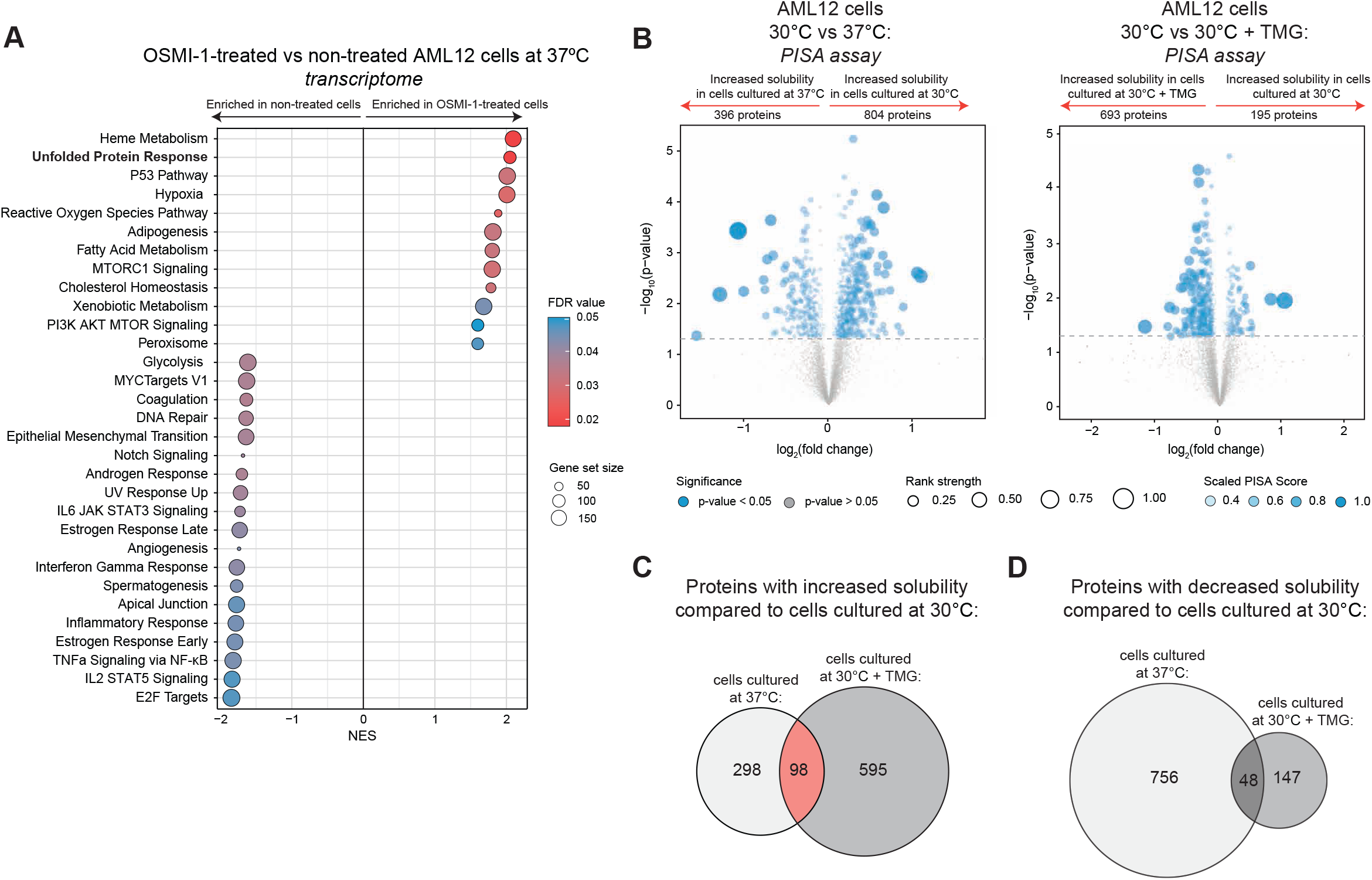
Transcriptome- and proteome-wide effects of O-GlcNAc manipulation in cultured cells. A) Gene Set Enrichment Analysis (GSEA) of Hallmark gene sets derived from RNA-seq data of AML12 cells treated with the OGT inhibitor OSMI-1 compared to untreated controls. Gene sets with an FDR q-value < 0.1 are shown. B) Volcano plot of PISA assay data showing differential thermal aggregation comparing cells cultured at 30 °C vs 37 °C and at 30 °C ± TMG. Point size reflects rank strength based on the composite PISA score (−log_10_(p-value) × |log_2_ fold change|), where a higher score corresponds to a better rank. Opacity reflects the PISA score scaled to 0.3-1 for visualization. C, D) Venn diagram showing the overlap of proteins in cells cultured at 37 °C or 30 °C + TMG with significantly increased (C) or decreased (D) solubility (*p* < 0.05) compared to cells cultured at 30 °C. Statistics: For GSEA analysis, enrichment significance was determined using normalized enrichment scores (NES) and false discovery rate (FDR q-values) calculated by the Broad Institute GSEA algorithm. Gene sets with FDR q-value < 0.1 were considered significantly enriched.

## Discussion

O-GlcNAcylation is an evolutionarily conserved protein monoglycosylation, and the glycosyltransferase, OGT, for this modification is an essential gene in at least *Drosophila*, mice and cultured mammalian cells (Hardivillé and Hart, 2014; Zachara et al., 2022). We found broad loss of protein O-GlcNAc in mitochondrial CIII-deficient mice and, initially, presumed that this reflects insufficient hexosamine biosynthesis. Hexosamine biosynthesis and O-GlcNAcylation are frequently described as nutrient sensors (Paneque et al., 2023; Yang and Qian, 2017), yet their activity in mitochondrial diseases, conditions often characterized by severely compromised energy metabolism remains largely unexplored. Unexpectedly, we found that UDP-GlcNAc levels are maintained whereas hypothermia leads to the loss of O-GlcNAc in mice with severe OXPHOS deficiency. Our data from the mutant mice and other models show an evolutionarily conserved relationship between temperature and global cellular O-GlcNAc status.

We previously showed that O-GlcNAcylation is not sensitive to changes in nutrient availability and HBP flux in cultured cells. In fact, it was affected only under extreme conditions, such as full loss of the HBP (Sunden et al., 2023). The lack of correlation between UDP-GlcNAc and O-GlcNAc levels in mice was consistent with these cell data. The resistance is further supported by the wide range of UDP-GlcNAc concentrations over which OGT remains catalytically active (Zachara et al., 2022). Our data also make OGT inhibition by UDP an unlikely scenario. Neither GlcNAc supplementation nor genetic enhancement of the HBP restored O-GlcNAc levels *in vivo*. Notably, neither intervention increased UDP-GlcNAc levels, in contrast to the robust increases we observed in cultured cells (Sunden et al., 2023). This likely reflects tight *in vivo* regulation of the HBP as GFPT1 is allosterically regulated by its products, including glucosamine-6-phosphate and UDP-GlcNAc (Jia et al., 2020). Together, these data ruled out impaired hexosamine biosynthesis as the driver of O-GlcNAc loss in OXPHOS-deficient mice despite their severe metabolic stress.

A study from 2014 reported that O-GlcNAc levels in *Drosophila melanogaster* embryos correlate with ambient temperature (Radermacher et al., 2014). We extended this study by showing that temperature similarly regulates O-GlcNAc levels in adult fruit flies, zebrafish and mammalian cells across physiological temperature ranges. Three independent mouse interventions that partially or fully corrected the body temperature of the hypothermic *Bcs1l^p.S78G^*mice also restored O-GlcNAc levels, demonstrating that body temperature was the key determinant of O-GlcNAc status in mice. To interrogate the functional significance of the O-GlcNAc loss, we restored O-GlcNAc levels in the mutant mice by inhibiting OGA. This intervention did not ameliorate disease phenotypes but instead even slightly exacerbated some, indicating that reduced O-GlcNAc levels are not inherently detrimental in hypothermia. Strikingly, elevating O-GlcNAc levels had minimal impact on the hepatic and skeletal muscle transcriptomes, strongly suggesting that decreased O-GlcNAc in hypothermia is a physiological response similar to that in poikilothermic animals, or, rather, that O-GlcNAcylation is evolutionarily adapted to the mammalian body temperature and less needed at lower temperatures. Mechanistically, hypothermia decreased the OGT/OGA ratio by decreasing OGT and increasing OGA transcription, favoring net O-GlcNAc removal. This regulation occurred independently of the previously described DI–mediated feedback pathway (Tan et al., 2020) indicating a distinct temperature-responsive mechanism. It remains to be studied which proteins or other molecules monitor the cellular temperature and which transcriptional regulators deliver this information to coordinate the OGT-OGA transcription with temperature.

O-GlcNAcylation has been implicated in the regulation of proteostasis and stress responses (Kazemi et al., 2010; Ryan et al., 2019; Wang et al., 2014; Wulff-Fuentes et al., 2023; Zeidan et al., 2024). Effects on protein stability, turnover, and degradation (Zachara et al., 2022) as well as co-translational stabilization of nascent proteins (Zeidan et al., 2024; Zhu et al., 2015) have been reported. Our PISA assay suggested that hypothermia induces bidirectional changes in protein solubility, with subsets of proteins displaying either increased or decreased solubility upon subsequent temperature elevation *in vitro*. In the fraction of proteins showing decreased solubility in cells cultured at 30 °C compared to 37 °C, 25% were similarly affected by OGA inhibition. In contrast, the reciprocal overlap was limited, as only 6% of proteins that were more soluble at 30 °C than at 37 °C also showed greater solubility at 30 °C than in TMG-treated cells. Compared to the large number of known O-GlcNAcylated proteins, we caught a solubility change in a relatively small number of proteins, possibly reflecting the technical limitations of the method. For example, nucleoporins that are some of the most highly O-GlcNAcylated cellular proteins are unlikely to be soluble under the conditions of the PISA assay. Recent chemical proteomics studies reported effects on the thermal solubility of a small fraction (72 proteins) of the proteome (King et al., 2022) or on biomolecular condensate formation (Xu et al., 2025). Together with our findings, these studies suggest that O-GlcNAc may influence the solubility, physical properties, or interactions of a subset of proteins rather than exerting a uniform effect across the proteome. Intriguingly, the reported effects are not unidirectional, suggesting that the relationship between O-GlcNAcylation and protein solubility may be context- and substrate-dependent. However, a uniform solubility-decreasing effect would be difficult to reconcile with the chemical properties of this neutral, hydrophilic modification. Consistent with this, a positive rather than negative relationship between protein solubility and O-GlcNAc levels would align with the general tendency of rising temperature to decrease protein solubility, as observed with some aggregation-prone proteins. Alternatively, O-GlcNAcylation may alter protein interactions or poorly soluble protein structures, such as biomolecular condensates. More detailed proteome-wide and protein-specific studies are required to confirm the solubility changes observed here and in the settings published previously.

While there is quite significant literature on the physiology of temperature adaptation of poikilothermic organisms, the literature is scarce on how individual proteins, protein complexes and the whole proteomes adapt to physiological temperature fluctuations (Guschina and Harwood, 2006; Li and Buck, 2023; Pucci and Rooman, 2017; Yamamoto et al., 2026). O-GlcNAcylation is evolutionarily conserved across eukaryotes, spanning organisms living in vastly different temperature optima and fluctuations. Interestingly, related forms of nucleocytoplasmic O-glycosylation are present even in systems lacking canonical OGT and OGA enzymes. In plants and protozoa, the OGT paralog SPINDLY (SPY) functions as a fucosyltransferase rather than an O-GlcNAc transferase (Mutanwad and Lucyshyn, 2022; Tiwari et al., 2025), while baker’s yeast (*Saccharomyces cerevisiae*) modifies nucleocytoplasmic proteins with O-mannose in a distribution similar to O-GlcNAc in multicellular organisms (Bartels et al., 2016). The recurrent use of neutral sugar moieties for nucleocytoplasmic protein O-monoglycosylation across evolution suggests a conserved functional role that is likely protome-wide rather than protein-specific. Because O-GlcNAcylation predates homeothermy in evolution, its original function may have been the protection of the proteome against environmental temperature fluctuations. Such function may be vital also in homeotherms to adjust to hibernation in some species, to hypothermia or hyperthermia due to adverse environmental conditions, and to the hyperthermia caused by inflammation.

The mechanisms by which O-GlcNAc exerts its biological functions remain largely unknown. No general binding motif or domain mediating O-GlcNAc-dependent intra- or intermolecular interactions have been identified in mammals. Taken together, our findings support a paradigm, in which O-GlcNAc acts as a global, evolutionarily conserved proteome adaptor to temperature fluctuations in both poikilothermic and homeothermic organisms. Our findings suggest that some of the diverse protein-specific effects attributed to O-GlcNAcylation may arise from general effects on protein stability or solubility.

## Materials and methods

### Mouse lines and husbandry

The *Bcs1l^p.S78G^;mt-Cyb^p.D254N^*, *Bcs1l^p.S78G^*and AOX mouse lines have been described previously (Levéen et al., 2011; Purhonen et al., 2020; Rajendran et al., 2019). The *Bcs1l^p.S78G^* mice on *mt-Cyb^p.D254N^* background (for simplicity referred to as *Bcs1l^p.S78G^*mice in the main text) were used as experimental model unless otherwise stated. *Bcs1l* WTs and heterozygote mice, irrespective of the *mt-Cyb^p.D254N^* variant, were considered phenotypically WT. The analyses included both sexes. All mice were on C57BL/6JCrl (RRID: IMSR_JAX:000664) nuclear genomic background and had *ad libitum* access to water and chow (Teklad 2018, Harlan). The mice were housed at the University of Helsinki animal facility in temperature-controlled conditions (23 °C), in individually ventilated cages under a 12-h light/dark cycle. All animal experiments were approved by the animal ethics committee of the State Provincial Office of Southern Finland (permit numbers ESAVI/6365/04.10.07/2017, ESAVI/16278/2020 and ESAVI/31141/2023) and conducted in accordance with FELASA (Federation of Laboratory Animal Science Associations) guidelines and 3R principles.

### Dietary interventions in mice

Powdered chow was supplemented with 1% (w/w) N-acetylglucosamine (Sigma-Aldrich, St. Louis, MO, USA) and 0.5% triacetyluridine (ChemCruz, Dallas, TX, USA). The chow was moistened, pressed into pellets, dried at room temperature, and stored at −20°C until use. The treated mice received supplemented chow *ad libitum* from weaning (P19-22) until sample collection.

### Thiamet G treatment in mice

Mice were treated with 20 mg/kg thiamet G (TMG, MedChemExpress, Monmouth Junction, NJ, USA), administered via daily intraperitoneal injection at P25-P29. Samples were collected at P29.

### Thermoneutrality intervention in mice

Mice were housed at 35 °C for three days with daily monitoring and sacrificed on P28.

### rAAV cloning, production and administration

For the cloning of the rAAV constructs, mouse *Bcs1l* and *Gfpt1* coding sequences were PCR-amplified and cloned into a pBluescript vector. Following sequencing, the fragments were subcloned into the hepatocyte-specific pAAV2-LSP1-EGFP vector (a kind gift from Prof. Ian Alexander, University of Sydney), containing the ApoE enhancer and human α1-antitrypsin promoter (Cunningham et al., 2015). The original EGFP-containing vector was used as control. Serotype 9 viral particles were produced by the AAV Gene Transfer and Cell Therapy Core Facility (University of Helsinki). The treated mice received 5×10^10^ viral particles in 100 µl saline via intraperitoneal injection at P21-22, and tissue samples were collected on P28 (Banerjee et al., 2026). GFPT1-V5 expression was verified by Western blot.

### Fly husbandry, strains and populations

*Drosophila melanogaster* strains were maintained at 25 °C under a 12-h light-dark cycle on standard medium. The *w^1118^* strain was used as wild-type control. The *oga^del.1^* strain (Akan et al., 2016) was provided by Dr. John Hanover (NIH, USA). For temperature treatments, five adult male flies (10-14 days old) were transferred to fresh vials and incubated at the indicated temperature for 16 h. *Drosophila montana*, originating from the populations Oulanka, Vancouver and Kamchatka were maintained at 19 °C under constant light on standard species-specific malt medium. For experiments, three adult male flies (21 days old) were transferred to fresh vials and incubated at the indicated temperature for 24 h.

### Zebrafish husbandry and strains

Zebrafish (AB strain) were housed at a density of 5 to 8 fish per tank in mixed-sex groups in 3-L tanks on a recirculating system (Aquatic Habitats) at 28.5 °C under a 14:10-h light-dark cycle. The fish were fed three times daily a commercial pelleted diet (Special Diet Services, SDS400) and Artemia nauplii (Sanders, Great Salt Lake Artemia). For the temperature treatments, five adult fish were transferred to water of indicated temperatures for 24 h.

### Sample collection

Mice were euthanized by cervical dislocation at P28-30 unless stated otherwise. Prior to collection, mice were fasted for two hours, and samples were collected during the light phase. Skin temperature was measured with the Gentle Temp 720 IR meter (Omron Health Care, Tokyo, Japan). Core temperature was measured by inserting a thermometer into the abdominal cavity (Testo 106, Testo SE & Co, Titisee-Neustadt, Germany). Blood glucose levels were measured using a Freestyle Lite meter (Abbott, Alameda, CA, USA) from blood within the body cavity during sample collection. Tissues were either fixed in 10% histology-grade formalin or snap frozen in liquid nitrogen and stored at −80 °C. Fly samples and zebrafish muscle tissue were snap-frozen and stored at −80 °C.

### Cell culture and treatments

AML12 cells (ATCC, #CRL-2254) were cultured in DMEM-F-12 medium with 10% fetal bovine serum, 5 mM glucose, glutamine, insulin-transferrin-selenium supplement (Gibco, Waltham, MA, USA), and penicillin/streptomycin. Hepa1-6 mouse hepatoma cells and human neonatal dermal fibroblast were cultured in DMEM with 10% fetal bovine serum, 5 mM glucose, glutamine, penicillin, and streptomycin. For the generation of the AML12 cells lines stably expressing tagged mouse OGT and OGA, the respective cDNAs were PCR-amplified and cloned in frame with an N-terminal mNeonGreen tag in the pcDNA6A vector. The mNeonGreen-OGT insert was subcloned into the pSB-BI-hyg and the mNeonGreen-OGA insert into pSB-puro. pSB-BI-hyg was a gift from Eric Kowarz (Addgene plasmid #60524; http://n2t.net/addgene:60524; RRID Addgene_60524). Cells were transfected with the OGT, OGA and the Sleeping Beauty transposon-encoding plasmids (Kowarz et al., 2015) using the FuGENE® HD Transfection Reagent according to the manufacturer’s instructions (Promega, Madison, Wisconsin, United States). Stable cell pools expressing mNeonGreen-OGT and mNeonGreen-OGA were selected using 2 μg/mL puromycin (Merck, P4512) and 500 μg/mL hygromycin B (Merck, H3274).

Cells were maintained at 37 °C and plated at least one day prior to experiments. At 40% confluency, cells were transferred to the indicated temperatures for 0-16 h. For OGA inhibition, AML12 and Hepa1-6 cells were treated with 0.5 μM TMG for 16 h at 30 °C or 37 °C. OGT was inhibited by treating AML12 cells 16 h with 50 µM OSMI-1 (MedChemExpress). After treatment, the cells were harvested in cold 80% MeOH, or in 2% SDS lysis buffer (60 mM Tris-Cl pH 6.8, 2% SDS, 12.5% glycerol, 1 mM EDTA) supplemented with protease inhibitors (Complete protease inhibitor mix, Roche, Basel, Switzerland), or RNAzol (Sigma-Aldrich) depending on the downstream analysis.

### SDS-PAGE and Western blot

Tissues (5-10-mg) or cells were homogenized in 2% SDS lysis buffer supplemented with protease inhibitors (Complete protease inhibitor mix, Roche) and briefly sonicated. Protein concentrations were determined using Bradford reagent (Bio-Rad, Hecules, CA, USA) with 2.5 mg/ml α-cyclodextrin (Sigma-Aldrich) to chelate interfering SDS.

For SDS-PAGE, bromophenol blue and 5% β-mercaptoethanol were added to the protein lysates and the samples were heated at 95°C for 5 minutes. Per lane, 5–30 µg total protein was run in TGX^TM^ precast gradient gels (Bio-Rad). Transfer onto PVDF (Amersham Hybond P 0.2 PVDF, Cytiva, Marlborough, MA) membranes was performed using the tank transfer method with Towbin buffer containing 10% MeOH. Equal loading and transfer were verified with Coomassie G-250.

Supplementary Table 3 lists all antibodies used. Detection and documentation of immunoblots were performed using peroxidase-conjugated secondary antibodies, enhanced chemiluminescence, and ChemiDoc MP (Bio-Rad) or Li-Cor Odyssey M (Lincoln, NE, USA) imaging systems. Prior to immunodetection of biotin-conjugated lectins, endogenous biotin was blocked with 1 µg/ml streptavidin (Jackson ImmunoResearch, West Grove, PA, USA), whereafter remaining biotin-binding sites were neutralized with 50 µM biotin (ChemCruz). All samples subjected to quantification were processed and run in a randomized order. For representative blots, samples from each experimental group were combined to generate an average signal, while all quantifications were performed on individually analyzed samples.

### RNA sequencing and differential expression analysis

Total RNA was extracted from cells or snap-frozen tissue using the RNAzol RT reagent (Sigma-Aldrich) following the manufacturer’s instructions and sent to Novogene (Beijing, China) for mRNA sequencing. Raw sequencing reads were processed by Novogene. Downstream analysis of the count data was performed in R (version 4.4.2) using the DESeq2 package (version 1.46.0). Variance stabilizing transformation (VST) normalized read counts, generated using DESeq2, were used to perform principal component analysis (PCA) for visualization of sample clustering and variability. In differential expression analysis, genes with an adjusted p-value of < 0.05 and an absolute log2 fold change > 1 were considered differentially expressed. Gene set enrichment analysis (GSEA) was performed using normalized gene expression values derived from DESeq2 with the Broad Institute GSEA software (v4.3.3). Samples were grouped according to experimental condition, and enrichment was assessed using the Hallmark gene set collection (MSigDB).

### Quantitative PCR

For qPCR analyses, total RNA was extracted from cells or snap-frozen tissue samples with RNAzol RT reagent (Sigma-Aldrich) according to manufacturer’s instructions. cDNA was synthesized, followed by qPCR using EvaGreen and Phire II Hot Start DNA polymerase-based detection chemistry, as described (Purhonen et al., 2020). The qPCR and data analysis were performed using a CFX96 thermocycler and CFX Manager software (Bio-Rad). PCR efficiencies were calculated using LinRegPCR software. The following primers were used to amplify cDNA:

*Ogt prod*: 5′– TGTGTTCGCAGTGACCT–3′, 5′–CAGCCAAATCTCCCCTTGTG–3′, *Ogt DI*: 5′– CCTGGGTCGCTTGGAAGAA–3′, 5′–CCCAGTCCTGCTCTAGTCTG–3′, *Ogt PTC*: 5′– CCTGGGTCGCTTGGAAGAA–3′, 5′–GACTTTTCGGTGCGTTCCTT–3′, *Oga prod*: 5′– TCGTAGTCACTCTTCAGCACA–3′, 5′–TCTTGTACACGGAAGCCTCA–3′, *Oga DI*: 5′– TCACAGCCTTGCCTTGAAAC–3′, 5′–TTGGAGGTTGGAGTCAGTGG–3′, *Oga PTC*: 5′– GTGCAGTGGTTAGCGTTTG–3′, 5′–GACGAAGCAGTAATCCAGGC–3′, *Rab11a*: 5′– AAGGCACAGATATGGGACACA–3′, 5’–CCTACTGCTCCACGATAGTATGC–3′, *Gak*: 5′-TTCCTCAACTGCCTTCCCATC-3′, 5′-ACGTATCCCATCCTCCTAGCA-498 3′. *Rab11a* and *Gak* were used as reference genes.

### Histology and cell counting

Formalin-fixed paraffin-embedded liver and kidney tissue sections were used for immunohistochemical staining of O-GlcNAc (antibody: O-GlcNAc, clone RL2, BioLegend, San Diego, CA, USA). For detection, Vectastain Elite ABC peroxidase or alkaline phosphatase reagents, or ImmPRESS peroxidase or alkaline phosphatase polymer detection reagents (Vector Laboratories, Newark, CA, USA) were used. Apoptotic cells were counted from H&E sections based on characteristic morphological features according to guidelines (Elmore et al., 2016).

### Metabolite analyses

UDP-HexNAc were quantified using UPLC-MS as described (Sunden et al., 2023). UDP-GlcNAc and uridine nucleotides were quantified using protocols previously developed and described by our laboratory (Purhonen et al., 2023b; Sunden et al., 2023; Upadhyay et al., 2024). For Ac-CoA quantification, ∼10 mg liver samples were homogenized in 0.4 ml ice-cold buffered 60% MeOH (10 mM Bis-Tris-Cl pH 7.0), containing 10 mM N-ethylmaleimide (NEM) to alkylate CoA. After pelleting insoluble material, the extracts were incubated for 5 minutes at 30 °C to complete the alkylation. Further precipitation of macromolecules and phase separation was induced by addition of 0.18 ml chloroform. The aqueous fractions were washed twice with 1.6 ml chloroform to remove MeOH and NEM. Any residual NEM was quenched with 1 mM DTT. Ac-CoA was quantified with coupled enzymatic reactions leading to a cyclic accumulation of NADH, the rate of which depended on the Ac-CoA + CoA concentration and served as a readout for the assay (increase in 340 nm absorbance). The assay reactions comprised: 100 mM Tris-Cl pH 7.5, 10 mM malate, 0.5 mg/ml fatty acid-free BSA, 1.9 mM acetyl phosphate, 1.1 mM NAD^+^, 1 U/ml malate dehydrogenase (Megazyme, Bray, Ireland), 1.9 U citrate synthase (Megazyme), and 14.4 U/ml phosphotransacetylase (Megazyme). The Gln/Glu ratio was calculated from a published metabolomics dataset (Kotarsky et al., 2012).

### OGT and OGA activity assays

The liver tissues were homogenized in PBS containing 0.1% Triton X-100, 2 mM β-mercaptoethanol, and protease inhibitor mix (EDTA-free Complete protease inhibitor mix, Roche) and clarified by centrifugation. OGA activity was determined by following the cleavage of a fluorogenic substrate 4-methylumbelliferyl N-acetyl-β-D-glucosaminide (4-MUF-NAG). The assay reactions comprised 2 mM 4-MUF-NAG, 0.5 mg/ml BSA, 2 mM β-mercaptoethanol, and 20 mM N-acetylgalactosamine in 50 mM potassium phosphate buffer pH 7.0. The inclusion of N-acetylgalactosamine served to inhibit lysosomal hexosaminidases. The background was determined in the presence of 25 µM TMG. OGT activity was determined from the same lysates (0.7 mg protein) after immunoprecipitation of OGT with paramagnetic protein A beads (Dynabeads, Invitrogen, Waltham, MA USA) and 500 ng antibody against OGT (Novus, NBP1-32791). On-bead O-GlcNAcylation reactions (10 µl) were carried out in 25 mM Bis-Tris-Cl pH 7.0, 0.3 mg/ml BSA, 2 mM β-mercaptoethanol, 100 µM ultra-pure UDP-GlcNAc (Promega, Madison, WI, USA), and 25 µg/ml substrate peptide (KKKYPGGSTPVSSANMM). After a 3 h incubation at room temperature (21-23°C), the reactions were stopped by separating the beads and the reaction media. OGT activities were determined based on the UDP released from UDP-GlcNAc. The released UDP was quantified using Promega UDP-Glo reagent. Background was determined in the presence of 36 µM OSMI-1. Purified immunoprecipitated recombinant OGT served as a positive control and normal rabbit serum (200 nl) immunoprecipitated lysate as a negative control. Due to the nature of the assay, lysates from 2-3 distinct mice of the same genotype were pooled together.

Activity of purified mouse OGA (50 ng/ml) as a function of temperature was measured using 4-MUF-NAG as the substrate as described for measurements from tissue samples, except that N-acetylgalactosamine was omitted and the reactions were carried out in a qPCR instrument (BioRad CFX384) with a temperature gradient. Full activity of the enzyme required 50 min pre-incubation at 37 °C before the experiment. The activity of purified human OGT (8.6 pM) on full-length protein substrates was measured using Promega UDP-Glo reagent. The O-GlcNAcylation reactions were carried out for 1 h in 25 mM Bis-Tris-Cl pH 7.0, 0.5 mg/ml BSA, 2 mM β-mercaptoethanol, and 200 µM ultra-pure UDP-GlcNAc. OGT activity was determined from accumulated UDP. The purified human full-length OGT was purchased from ActivMotif and caseine kinase 2 complex from New England Biolabs. Bovine α-crystallin derived from a Click-iT O-GlcNAc labeling kit (Thermo Scientific). His-tagged mouse OGA and its catalytically inactive D175N mutant version were produced in-house in E. coli BL21 and purified using metal affinity chromatography (HisPur Cobalt Resin, Thermo Scientific).

### Integrated PISA assay and Expression Proteomics (PISA-Express)

The proteome integral solubility alteration (PISA) assay and expression proteomics were carried out in an integrated way (PISA-Express) and analyzed to define difference in total abundance and solubility, where the changes of the latter was normalized to the total protein abundance change of each condition to correct for any changes in protein expression. All samples were combined into one TMT-based 18-plex experiment hosting both total abundance and solubility assays in one LC-MS/MS analysis, with a total of three conditions in triplicate for PISA and for expression proteomics, where these sample types originated from the same biological replicate either with or without temperature treatment gradient treatment and ultracentrifugation. AML12 cells were incubated at 30 °C or 37 °C with or without 0.5 μM TMG for 16 h. For each condition half sample was extracted as explained below and the other half was subjected to PISA.

### PISA sample processing

Samples were processed according to the proof-of-principle PISA method published article (Gaetani et al., 2019) and the latest optimized protocols for higher proteome depth of analysis, extended TMT multiplicity and improved sensitivity (Gaetani and Zubarev, 2023; Zhang et al., 2022). Briefly, cells were detached and washed twice with PBS, then resuspended in PBS supplemented with Halt™ protease inhibitor cocktail (Thermo Fisher Scientific). Cell lysis was performed using 0.4% NP-40 with five freeze–thaw cycles between liquid nitrogen and 37 °C. Lysates were clarified by ultracentrifugation, and the resulting soluble protein fraction was collected following application of the protein precipitation gradient. Samples from different temperature points were pooled prior to downstream analysis. For expression proteomics, samples were extracted without thermal treatment as described above and debris was removed by centrifugation at 14,000 at 4 °C for 30 min.

The total protein concentration of extracted samples from PISA and expression proteomics was measured using micro-BCA (bicinchoninic acid) assay. Briefly, as previously published in the above-mentioned references, 50 µg of each sample was reduced, alkylated and precipitated using cold acetone. Samples were resuspended in EPPS (4-(2-Hydroxyethyl)-1-piperazinepropanesulfonic acid) buffer, 8 M urea, pH 8.0, diluted down to 4 M urea and digested by LysC, then diluted down to 1 M urea and digested with trypsin. Each digest was labeled using TMTpro 18-plex technology (Thermo Fischer) and a final multiplex sample was first cleaned by Sep-Pack C18 column (Waters). The final sample was fractionated off-line by capillary reversed phase chromatography at pH 10 into 48 fractions, and each of them was then analyzed by high-resolution nLC–ESI-MS/MS (nanoscale liquid chromatography-electrospray ionization-tandem mass spectrometry), as described below. Then LC-MS/MS data were subjected to peptide / protein identification and quantification, and data analysis.

### PISA nanoLC-MS/MS analysis

NanoLC-MS/MS analyses were performed on an Orbitrap Exploris 480 mass spectrometer (Thermo Fisher Scientific). The instrument was equipped with an EASY ElectroSpray source and connected online to an Ultimate 3000 nanoflow UPLC system. The samples were pre-concentrated and desalted online using a PepMap C18 nano-trap column (length - 2 cm; inner diameter - 75 µm; particle size - 3 µm; pore size - 100 Å; Thermo Fisher Scientific) with a flow rate of 3 µL/min for 5 min. Peptide separation was performed on an EASY-Spray C18 reversed-phase nano-LC column (Acclaim PepMap RSLC; length - 50 cm; inner diameter - 2 µm; particle size - 2 µm; pore size – 100 Å; Thermo Scientific) at 55 °C and a flow rate of 300 nL/min. Peptides were separated using a binary solvent system consisting of 0.1% (v/v) FA, 2% (v/v) ACN (solvent A) and 98% ACN (v/v), 0.1% (v/v) FA (solvent B). They were eluted with a gradient of 3–26% B in 97 min, and 26–95% B in 9 min. Subsequently, the analytical column was washed with 95% B for 5 min before re-equilibration with 3% B. The mass spectrometer was operated in a data-dependent acquisition mode. A survey mass spectrum (from m/z 375 to 1500) was acquired in the Orbitrap analyzer at a nominal resolution of 120,000. The automatic gain control (AGC) target was set as 100% standard, with the maximum injection time of 50 ms. The most abundant ions in charge states 2+ to 7+ were isolated in a 3 s cycle, fragmented using HCD MS/MS with 33% normalized collision energy, and detected in the Orbitrap analyzer at a nominal mass resolution of 50,000. The AGC target for MS/MS was set as 250% standard with a maximum injection time of 100 ms, whereas dynamic exclusion was set to 45 s with a 10-ppm mass window.

### PISA protein identification, quantification and data analysis

Proteome Discoverer 3.1 was utilized for the database search and quantification against the UniProt Mouse protein database UP000000589. Cysteine carbamidomethylation was set as a fixed modification, along with TMT-related modifications, deamination on methionine, oxidation on arginine, and asparagine as variable modifications. Enzyme specificity was defined as trypsin with a maximum of two missed cleavages. A 1% false discovery rate was employed as a filter at both the protein and peptide levels. Contaminants and reversed-hit peptides were removed, and only proteins with at least two unique peptides were included in the quantitative analysis. Proteins with missing values in all replicates for any treatment were eliminated. The quantified abundance of each protein in each sample (labeled with a different TMT) was normalized to the total intensity of all proteins in that sample.

### Statistics

All samples were collected, processed, and analyzed in a randomized order. In the figures, error bars represent the mean and 95% CI of the mean. For comparisons between two groups, differences were analyzed using Welch’s t-test, or, when normality assumptions were not met, the Mann–Whitney U test. For comparisons involving more than two groups, one-way ANOVA followed by pre-selected pairwise comparisons (Welch’s t-tests) were used. For experiments with two independent factors, data were analyzed using two-way ANOVA followed by Šídák’s multiple comparisons test. For data normalized to internal controls and expressed as fold change relative to control, statistical significance was assessed using a one-sample t-test. All pairwise comparisons were performed using two-sided tests. In mouse and zebrafish experiments, each experimental unit (*n*) corresponds to an individual animal, except for the experiment regarding OGT activity, in which pooled lysates from individual mice were used. In *Drosophila* experiments, each experimental unit consisted of 3-5 pooled flies. Within each experiment, at least three independent experimental units originating from different fly generations were used. Cell culture experiments were performed in three parallel replicates per condition within each experiment. GraphPad Prism software was used for statistical analyses (GraphPad Software Inc., La Jolla, CA, USA). The figure legends provide information on the specific statistical tests applied and the *p*-values are displayed in the graphs.

## Acknowledgments

We thank Vilma Wanne for technical assistance. We thank the core facilities of University of Helsinki, Nina Sipari of the Viikki Metabolomics Unit, the Tissue Preparation and Histochemistry Unit (Department of Anatomy) and the Finnish Centre for Laboratory Animal Pathology (Faculty of Veterinary Medicine) for processing of histological samples, the Laboratory Animal Center of the University of Helsinki for the animal husbandry, and the Helsinki Fly Facility for provision of fly food. We thank Dr. Henri Koivula at the NC-ZebrafU, Helsinki In Vivo Animal Imaging Unit, University of Helsinki for the zebrafish sample collection. We acknowledge Chemical Proteomics at the Karolinska Institutet (Headed by Massimiliano Gaetani at Chemistry I Division, MBB Department), also Unit of SciLifeLab and node of the Swedish National Infrastructure for Biological Mass Spectrometry (BioMS), which provided full support in the experimental design, performance, and data analysis of the proteomic studies. We acknowledge funding from Folkhälsan Research Center, Jane and Aatos Erkko Foundation, the Foundation for Pediatric Research, Finska Läkaresällskapet, Medicinska Understödsföreningen Liv och Hälsa rf, Magnus Ehrnrooth Foundation, Sigrid Jusélius Foundation and University of Helsinki (The Doctoral Programme in Clinical Research). Graphical illustrations were created with BioRender (https://www.biorender.com/).

## Author contributions

JK, JP and CK designed and supervised the study. DU, CK, RB, JP and JK performed mouse experiments and sample collection. DU, CK, RB, NS, JP and JK performed the mouse tissue analyses. CK performed the *Drosophila,* and zebrafish experiments and analyzed their data. MK was responsible for the *Drosophila montana* maintenance and sample collection. CK, DU and NS performed the cell culture experiments and analyzed their data. CK did the RNA seq transcriptomics analyses and analyzed the histological data. JP performed the OGT and OGA activity measurements. MG provided design, performance and bioinformatic data analysis of the PISA assay. CK was responsible for the statistics. DU, CK and JK wrote the first manuscript draft. CK and DU prepared the figure panels. VF and all other authors critically read and commented on the manuscript, and JK and CK revised it accordingly.

## Data availability

All raw data, including transcriptomics and proteomics, is available from the authors upon request.

---

**Supplementary Figure 1.**

A) Representative O-GlcNAc immunohistochemistry of kidney sections of WT and *Bcs1l^p.S78G^* (*mt-Cyb^p.D254N^* background) mice.

B-D) Representative lectin blots with wheat germ agglutinin (WGA) and concanavalin A (Con A) in liver (B), kidney (C), and skeletal muscle (D) lysates from WT and *Bcs1l^p.S78G^;mt-Cyb^p.D254N^* mice, n = 5-6/genotype.

E-G) Representative western blot and quantification of O-GlcNAc levels in the liver (E), kidney (F), and skeletal muscle (G) lysates of WT and *Bcs1l^p.S78G^* mice on WT mtDNA background at P200, n = 8/genotype.

H) Concentration of hepatic UDP-HexNAc n = 8/genotype, from WT and *Bcs1l^p.S78G^*mice (*mt-Cyb^p.D254N^* background).

Statistics: Welch’s two-sided t-test. *p* values of the group comparisons are indicated above the panel. The error bars represent 95% CI of mean. All data points derive from independent mice.

**Supplementary Figure 2.**

A) Representative western blot and quantification of O-GlcNAc levels in Hepa1-6 cells incubated 16 h at indicated temperatures, n = 3/condition.

B) Representative western blot and quantification of O-GlcNAc levels in Hepa1-6 cells maintained for the indicated time points at 30°C, n = 3/condition.

C) Representative lectin blots with WGA and Con A in AML12 cells incubated 16 h at indicated temperatures, n = 3/condition.

Statistics: One sample t-test. *p* values of the group comparisons are indicated above the panel. Error bars represent 95% CI of mean. The data derive from three independent experiments.

**Supplementary Figure 3.**

A) Relative activity of purified OGT on different O-GlcNAc acceptor proteins (OGA, OGA^D172N^, α-Crystallin and Casein kinase II) measured at different temperatures.

B) Activity of purified OGA as measured using the fluorogenic substrate 4-MUF-NAG across different temperatures with or without prewarming at 37 °C prior to measurement. TMG was used as negative control.

C) Representative western blot and quantification of OGT and OGA protein levels, and their ratio, in Hepa1-6 cells incubated 16 h at indicated temperatures, n = 3/condition.

D-F) Representative western blot and quantification of OGT and OGA protein levels, and their ratio, in skeletal muscle (D), kidney (E), and liver (F) tissues of WT and *Bcs1l^p.S78G^* mice on WT mtDNA background at P200, n = 7-8/genotype.

G) Correlation between hepatic OGT/OGA ratio (western blot quantification) and core temperature of WT and *Bcs1l^p.S78G^* mice on WT mtDNA background at P33.

Statistics: One sample t-test (C) and Welch’s two-sided t-test (D-F). *p* values of the group comparisons are indicated above the panel. Error bars represent 95% CI of mean. The data in C derive from three independent experiments. Data points in D-G derive from independent mice.

**Supplementary Figure 4.**

A) Representative western blot and quantification of O-GlcNAc, OGT and OGA protein levels in Hepa1-6 cells ± 0.5 µM thiamet G (TMG) incubated 16 h at 30 °C or 37 °C, n = 3.

B) Volcano plots of differentially expressed transcripts in AML12 cells cultured 16 h at 30 °C with or without TMG. Genes with |log2 fold change| > 1 and FDR adjusted p-value < 0.05 are considered significant.

C) Volcano plot of quantitative proteomics data from AML12 cells cultured 16 h at 30 °C with or without TMG. Proteins with |log2 fold change| > 0.5 and p-value < 0.05 are considered significant.

D) Representative western blot and quantification of O-GlcNAc levels in AML12 cells treated with 50 µM OSMI-1 for 16 h, n = 3.

E) Volcano plot of differentially expressed transcripts in AML12 cells treated with OSMI-1 compared to controls. Genes with |log2 fold change| > 1 and adjusted p-value < 0.05 are considered significant.

Statistics: Two-way ANOVA followed by Šídák’s multiple comparisons test for selected pairwise comparisons (A), and Welch’s two-sided t-test (D). Data in A and D derive from three independent experiments. For RNA-seq analyses (B and E), differential expression was determined using DESeq2 with multiple testing correction (Benjamini–Hochberg). For proteomics data (C), statistical significance was assessed using two-sided Welch’s t-test.

## References

Akan, I., Love, D.C., Harwood, K.R., Bond, M.R., Hanover, J.A., 2016. *Drosophila* O-GlcNAcase deletion globally perturbs chromatin O-GlcNAcylation. J Biol Chem 291, 9906–9919.

Akella, N.M., Ciraku, L., Reginato, M.J., 2019. Fueling the fire: emerging role of the hexosamine biosynthetic pathway in cancer. BMC Biol. 17, 52.

Banerjee, R., Purhonen, J., Kallijärvi, J., 2021. The mitochondrial coenzyme Q junction and complex III: biochemistry and pathophysiology. FEBS J.

Banerjee, R., Purhonen, J., Sultana, N., Ros, O., Nieminen, A.I., Kietz, C., Fellman, V., Kallijärvi, J., 2026. Hepatic gene replacement improves energy metabolism and survival in a mouse model of neonatal mitochondrial disease GRACILE syndrome. Mol Ther 34, 4518–4530.

Banerjee, R., Upadhyay, D., Zarybnický, T., Kietz, C., Kuure, S., Fellman, V., Purhonen, J., Kallijärvi, J., 2024. Hepatic mitochondrial respiration is crucial for euthermia in complex III-deficient mice with impaired brown adipose tissue thermogenesis. 10.1101/2024.09.23.612616

Bartels, M.F., Winterhalter, P.R., Yu, J., Liu, Y., Lommel, M., Möhrlen, F., Hu, H., Feizi, T., Westerlind, U., Ruppert, T., Strahl, S., 2016. Protein O-mannosylation in the murine brain: occurrence of mono-O-mannosyl glycans and identification of new substrates. PLoS One 11, e0166119.

Bornstein, R., Mulholland, M.T., Sedensky, M., Morgan, P., Johnson, S.C., 2023. Glutamine metabolism in diseases associated with mitochondrial dysfunction. Mol Cell Neurosci 126, 103887.

Cunningham, S.C., Siew, S.M., Hallwirth, C.V., Bolitho, C., Sasaki, N., Garg, G., Michael, I.P., Hetherington, N.A., Carpenter, K., de Alencastro, G., Nagy, A., Alexander, I.E., 2015. Modeling correction of severe urea cycle defects in the growing murine liver using a hybrid recombinant adeno-associated virus/piggyBac transposase gene delivery system. Hepatology 62, 417–428.

El-Khoury, R., Rak, M., Bénit, P., Jacobs, H.T., Rustin, P., 2022. Cyanide resistant respiration and the alternative oxidase pathway: A journey from plants to mammals. Biochim Biophys Acta Bioenerg 1863, 148567.

Elmore, S.A., Dixon, D., Hailey, J.R., Harada, T., Herbert, R.A., Maronpot, R.R., Nolte, T., Rehg, J.E., Rittinghausen, S., Rosol, T.J., Satoh, H., Vidal, J.D., Willard-Mack, C.L., Creasy, D.M., 2016. Recommendations from the INHAND apoptosis/necrosis working group. Toxicol Pathol 44, 173–188.

Fellman, V., Rapola, J., Pihko, H., Varilo, T., Raivio, K.O., 1998. Iron-overload disease in infants involving fetal growth retardation, lactic acidosis, liver haemosiderosis, and aminoaciduria. Lancet 351, 490–493.

Gaetani, M., Sabatier, P., Saei, A.A., Beusch, C.M., Yang, Z., Lundström, S.L., Zubarev, R.A., 2019. Proteome Integral Solubility Alteration: A high-throughput proteomics assay for target deconvolution. J Proteome Res 18, 4027–4037.

Gaetani, M., Zubarev, R.A., 2023. Proteome Integral Solubility Alteration (PISA) for high-throughput ligand target deconvolution with increased statistical significance and reduced sample amount. Methods Mol Biol 2554, 91–106

Guschina, I.A., Harwood, J.L., 2006. Mechanisms of temperature adaptation in poikilotherms. FEBS Letters 580, 5477–5483.

Hardivillé, S., Hart, G.W., 2014. Nutrient regulation of signaling, transcription, and cell physiology by O-GlcNAcylation. Cell Metab 20, 208–213.

Hoikkala, A., Poikela, N., 2022. Adaptation and ecological speciation in seasonally varying environments at high latitudes: *Drosophila virilis* group. Fly 16, 85–104.

Holt, G.D., Hart, G.W., 1986. The subcellular distribution of terminal N-acetylglucosamine moieties. Localization of a novel protein-saccharide linkage, O-linked GlcNAc. J Biol Chem 261, 8049–8057.

Issad, T., Al-Mukh, H., Bouaboud, A., Pagesy, P., 2022. Protein O-GlcNAcylation and the regulation of energy homeostasis: lessons from knock-out mouse models. J Biomed Sci 29, 64.

Jia, C., Li, H., Fu, D., Lan, Y., 2020. GFAT1/HBP/O-GlcNAcylation axis regulates β-catenin activity to promote pancreatic cancer aggressiveness. Biomed Res Int 2020, 1921609.

Kazemi, Z., Chang, H., Haserodt, S., McKen, C., Zachara, N.E., 2010. *O*-Linked β-*N*-acetylglucosamine (*O*-GlcNAc) regulates stress-induced heat shock protein expression in a GSK-3β-dependent manner. Journal of Biological Chemistry 285, 39096–39107.

King, D.T., Serrano-Negrón, J.E., Zhu, Y., Moore, C.L., Shoulders, M.D., Foster, L.J., Vocadlo, D.J., 2022. Thermal proteome profiling reveals the O-GlcNAc-dependent meltome. J Am Chem Soc 144, 3833–3842.

Kotarsky, H., Keller, M., Davoudi, M., Levéen, P., Karikoski, R., Enot, D.P., Fellman, V., 2012. Metabolite profiles reveal energy failure and impaired beta-oxidation in liver of mice with complex III deficiency due to a BCS1L mutation. PLoS ONE 7, e41156.

Kowarz, E., Löscher, D., Marschalek, R., 2015. Optimized Sleeping Beauty transposons rapidly generate stable transgenic cell lines. Biotechnol J 10, 647–653.

Levéen, P., Kotarsky, H., Mörgelin, M., Karikoski, R., Elmér, E., Fellman, V., 2011. The GRACILE mutation introduced into *Bcs1l* causes postnatal complex III deficiency: a viable mouse model for mitochondrial hepatopathy. Hepatology 53, 437–447.

Li, Z.-L., Buck, M., 2023. A proteome-scale analysis of vertebrate protein amino acid occurrence: Thermoadaptation shows a correlation with protein solvation but less so with dynamics. Proteins 91, 3–15. 10.1002/prot.26404

Martinez, M.R., Dias, T.B., Natov, P.S., Zachara, N.E., 2017. Stress-induced O-GlcNAcylation, an adaptive process of injured cells. Biochem Soc Trans 45, 237–249.

Mutanwad, K.V., Lucyshyn, D., 2022. Balancing O-GlcNAc and O-fucose in plants. FEBS J 289, 3086–3092.

Paneque, A., Fortus, H., Zheng, J., Werlen, G., Jacinto, E., 2023. The hexosamine biosynthesis pathway: regulation and function. Genes 14, 933.

Pucci, F., Rooman, M., 2017. Physical and molecular bases of protein thermal stability and cold adaptation. Curr Opin Struct Biol 42, 117–128.

Purhonen, J., Banerjee, R., Wanne, V., Sipari, N., Mörgelin, M., Fellman, V., Kallijärvi, J., 2023a. Mitochondrial complex III deficiency drives c-MYC overexpression and illicit cell cycle entry leading to senescence and segmental progeria. Nat Commun 14, 2356.

Purhonen, J., Grigorjev, V., Ekiert, R., Aho, N., Rajendran, J., Pietras, R., Truvé, K., Wikström, M., Sharma, V., Osyczka, A., Fellman, V., Kallijärvi, J., 2020. A spontaneous mitonuclear epistasis converging on Rieske Fe-S protein exacerbates complex III deficiency in mice. Nat Commun 11, 322.

Purhonen, J., Hofer, A., Kallijärvi, J., 2023b. Quantification of all 12 canonical ribonucleotides by real-time fluorogenic *in vitro* transcription. Nucleic Acids Research gkad1091.

Radermacher, P.T., Myachina, F., Bosshardt, F., Pandey, R., Mariappa, D., Müller, H.-A.J., Lehner, C.F., 2014. O-GlcNAc reports ambient temperature and confers heat resistance on ectotherm development. PNAS 111, 5592–5597.

Rajendran, J., Purhonen, J., Tegelberg, S., Smolander, O.-P., Mörgelin, M., Rozman, J., Gailus-Durner, V., Fuchs, H., Hrabe de Angelis, M., Auvinen, P., Mervaala, E., Jacobs, H.T., Szibor, M., Fellman, V., Kallijärvi, J., 2019. Alternative oxidase-mediated respiration prevents lethal mitochondrial cardiomyopathy. EMBO Mol Med 11.

Ryan, P., Xu, M., Davey, A.K., Danon, J.J., Mellick, G.D., Kassiou, M., Rudrawar, S., 2019. O-GlcNAc modification protects against protein misfolding and aggregation in neurodegenerative disease. ACS Chem. Neurosci. 10, 2209–2221.

Slawson, C., Zachara, N.E., Vosseller, K., Cheung, W.D., Lane, M.D., Hart, G.W., 2005. Perturbations in O-linked beta-N-acetylglucosamine protein modification cause severe defects in mitotic progression and cytokinesis. J Biol Chem 280, 32944–32956.

Sunden, M., Upadhyay, D., Banerjee, R., Sipari, N., Fellman, V., Kallijärvi, J., Purhonen, J., 2023. Enzymatic assay for UDP-GlcNAc and its application in the parallel assessment of substrate availability and protein O-GlcNAcylation. Cell Rep Methods 3, 100518.

Tan, Z.-W., Fei, G., Paulo, J.A., Bellaousov, S., Martin, S.E.S., Duveau, D.Y., Thomas, C.J., Gygi, S.P., Boutz, P.L., Walker, S., 2020. O-GlcNAc regulates gene expression by controlling detained intron splicing. Nucleic Acids Res 48, 5656–5669.

Tiwari, M., Gas-Pascual, E., Goyal, M., Popov, M., Matsumoto, K., Grafe, M., Gräf, R., Haltiwanger, R.S., Olszewski, N., Orlando, R., Samuelson, J.C., West, C.M., 2025. Novel antibodies detect nucleocytoplasmic O-fucose in protist pathogens, cellular slime molds, and plants. mSphere 10, e0094524.

Toleman, C.A., Schumacher, M.A., Yu, S.-H., Zeng, W., Cox, N.J., Smith, T.J., Soderblom, E.J., Wands, A.M., Kohler, J.J., Boyce, M., 2018. Structural basis of O-GlcNAc recognition by mammalian 14-3-3 proteins. Proc Natl Acad Sci U S A 115, 5956–5961.

Torres, C.R., Hart, G.W., 1984. Topography and polypeptide distribution of terminal N-acetylglucosamine residues on the surfaces of intact lymphocytes. Evidence for O-linked GlcNAc. J Biol Chem 259, 3308–3317.

Upadhyay, D., Banerjee, R., Kallijärvi, J., Purhonen, J., 2024. Protocol for quantification of UDP-GlcNAc using an enzymatic microplate assay. STAR Protoc 5, 102817.

Visapää, I., Fellman, V., Vesa, J., Dasvarma, A., Hutton, J.L., Kumar, V., Payne, G.S., Makarow, M., Van Coster, R., Taylor, R.W., Turnbull, D.M., Suomalainen, A., Peltonen, L., 2002. GRACILE syndrome, a lethal metabolic disorder with iron overload, is caused by a point mutation in *BCS1L*. Am. J. Hum. Genet. 71, 863–876.

Wang, Z.V., Deng, Y., Gao, N., Pedrozo, Z., Li, D.L., Morales, C.R., Criollo, A., Luo, X., Tan, W., Jiang, N., Lehrman, M.A., Rothermel, B.A., Lee, A.-H., Lavandero, S., Mammen, P.P.A., Ferdous, A., Gillette, T.G., Scherer, P.E., Hill, J.A., 2014. Spliced X-box binding protein 1 couples the unfolded protein response to hexosamine biosynthetic pathway. Cell 156, 1179–1192.

Wulff-Fuentes, E., Boakye, J., Kroenke, K., Berendt, R.R., Martinez-Morant, C., Pereckas, M., Hanover, J.A., Olivier-Van Stichelen, S., 2023. O-GlcNAcylation regulates OTX2’s proteostasis. iScience 26, 108184.

Xu, S., Yin, K., Xu, X., Fu, L., Wu, R., 2025. O-GlcNAcylation reduces proteome solubility and regulates the formation of biomolecular condensates in human cells. Nat Commun 16, 4068.

Yamamoto, T., Shiraishi, A., Sakai, T., Wada, A., Satake, H., 2026. Hydrophobic interactions determine the optimum temperature of a housekeeping enzyme. Journal of Biological Chemistry 302, 111234.

Yang, X., Qian, K., 2017. Protein O-GlcNAcylation: emerging mechanisms and functions. Nat Rev Mol Cell Biol 18, 452–465.

Zachara, N.E., Akimoto, Y., Boyce, M., Hart, G.W., 2022. The O-GlcNAc Modification, in: Varki, A., Cummings, R.D., Esko, J.D., Stanley, P., Hart, G.W., Aebi, M., Mohnen, D., Kinoshita, T., Packer, N.H., Prestegard, J.H., Schnaar, R.L., Seeberger, P.H. (Eds.), Essentials of Glycobiology. Cold Spring Harbor Laboratory Press, Cold Spring Harbor (NY).

Zeidan, Q., Tian, J.L., Ma, J., Eslami, F., Hart, G.W., 2024. O-GlcNAcylation of ribosome-associated proteins is concomitant with translational reprogramming during proteotoxic stress. Journal of Biological Chemistry 300.

Zhang, X., Lytovchenko, O., Lundström, S.L., Zubarev, R.A., Gaetani, M., 2022. Proteome Integral Solubility Alteration (PISA) Assay in mammalian cells for deep, high-confidence, and high-throughput target deconvolution. Bio Protoc 12, e4556.

Zhang, Z., Tan, E.P., VandenHull, N.J., Peterson, K.R., Slawson, C., 2014. O-GlcNAcase expression is sensitive to changes in O-GlcNAc homeostasis. Front Endocrinol (Lausanne) 5, 206.

Zhao, Y., Li, R., Wang, W., Zhang, H., Zhang, Q., Jiang, J., Wang, Y., Li, Y., Guan, F., Nie, Y., 2024. O-GlcNAc signaling: Implications for stress-induced adaptive response pathway in the tumor microenvironment. Cancer Letters 598, 217101.

Zhu, Y., Liu, T.-W., Cecioni, S., Eskandari, R., Zandberg, W.F., Vocadlo, D.J., 2015. O-GlcNAc occurs cotranslationally to stabilize nascent polypeptide chains. Nat Chem Biol 11, 319–325.

