## Supplemental Table 3 for "Evolutionarily conserved temperature dependence leads to loss of protein O-GlcNAc in mammalian hypothermia"

**Supplementary Table 3**. List of antibodies used during western blotting and IHC.

| Target | Antibody type | Identifier | Clone | Source | Dilution | Blocking solution |
|---|---|---|---|---|---|---|
| O-GlcNAc | Mouse monoclonal | 677902 | RL2 | BioLegend | WB: 1:1000<br>IHC: 1:1000 | 0.7% Na-Caseinate-1% BSA |
| O-GlcNAc | Mouse monoclonal | 838004 | CTD110.6 | BioLegend | WB: 1:2000 | 0.7% Na-Caseinate-1% BSA |
| MGEA5 (OGA) | Rabbit polyclonal | HPA036141 | — | Atlas Antibodies | WB: 1:1000 | 5% fat-free milk-TBST |
| MGEA5 (OGA) | Rabbit monoclonal | 60406 | E9C5U | CST | WB: 1:2000 | 5% fat-free milk-TBST |
| OGT | Rabbit polyclonal | ab96718 | — | Abcam | WB: 1:4000 | 5% fat-free milk-TBST |
| mNeonGreen | Rabbit polyclonal | 29523-1-AP | — | Proteintech | WB: 1:6000 | 5% fat-free milk-TBST |
| Concanavalin A – biotinylated | Lectin (Biotin) | Biotinylated in-house | | | WB: 1:4000<br>1 mM $Mg^{2+}$ and $Ca^{2+}$ added | 1% PVP-0.5% PVA-TBST |
| WGA – biotinylated | Lectin (Biotin) | B-1025 | — | Vector laboratories | WB: 1:10 000<br>1 mM $Mg^{2+}$ and $Ca^{2+}$ added | 1% PVP-0.5% PVA-TBST |
| Anti-Rabbit IgG, HRP | Goat polyclonal (HRP) | 7074 | — | CST | WB: 1:2000–1:5000 | — |
| Anti-Mouse IgG, HRP | Goat polyclonal (HRP) | 7076 | — | CST | WB: 1:2000–1:5000 | — |
| Streptavidin-POD conjugate | — | 1108915-3001 | — | Roche | WB: 1:20 000 | — |
