## Supplemental Table 2 for "Evolutionarily conserved temperature dependence leads to loss of protein O-GlcNAc in mammalian hypothermia"

**Supplementary Table 2**. Proteins with significantly increased solubility ($log_2FC < 0$, $p < 0.05$) identified by PISA assay in cells cultured at 30°C vs 37°C and at 30°C vs 30°C + TMG.

| | **30°C vs 37°C** | | | | **30°C vs 30°C ± TMG** | | | | | |
|---|---|---|---|---|---|---|---|---|---|---|
| **Protein** | **$Log_2$ FC** | **p value** | **Score** | **Rank** | **$Log_2$ FC** | **p value** | **Score** | **Rank** | **Function** | **Functional group** |
| PLEC | 0.463 | 0.0002 | 1.677 | 21 | 0.323 | 0.023 | 0.527 | 186 | Cytolinker, scaffold protein | Cytoskeleton, membrane organization and trafficking |
| CFAP20 | 0.25 | 0.007 | 0.538 | 302 | 0.211 | 0.011 | 0.41 | 330 | Axonemal structure factor | |
| TJP1 | 0.223 | 0.016 | 0.401 | 505 | 0.213 | 0.023 | 0.347 | 430 | Tight junction scaffold protein | |
| SPTBN1 | 0.18 | 0.045 | 0.241 | 879 | 0.336 | 0.035 | 0.746 | 82 | Spectrin scaffold protein | |
| PFN1 | 0.175 | 0.046 | 0.234 | 895 | 0.314 | 0.038 | 0.672 | 117 | Actin polymerization regulator | |
| MAP1LC3A | 0.122 | 0.017 | 0.214 | 940 | 0.219 | 0.002 | 0.572 | 155 | Autophagy and autophagosome assembly factor | |
| ACTR10 | 0.139 | 0.049 | 0.181 | 1039 | 0.218 | 0.039 | 0.387 | 366 | Actin-related motor protein | |
| NSFL1C | 0.069 | 0.021 | 0.116 | 1163 | 0.111 | 0.009 | 0.228 | 677 | Vesicle fusion ATPase regulator | |
| CMAS | 0.276 | 0.014 | 0.514 | 333 | 0.239 | 0.038 | 0.341 | 441 | Sialic acid biosynthesis enzyme | Metabolism and small-molecule processing |
| CHLSN | 0.269 | 0.04 | 0.374 | 552 | 0.42 | 0.033 | 0.793 | 65 | Secreted cholesterol-regulating protein | |
| DBT | 0.116 | 0.004 | 0.275 | 764 | 0.16 | 0.001 | 0.459 | 257 | Branched-chain amino acid kinase | |
| ENOPH1 | 0.16 | 0.032 | 0.238 | 886 | 0.135 | 0.040 | 0.189 | 783 | Metabolic enzyme enolase-like protein | |
| NANS | 0.097 | 0.034 | 0.143 | 1120 | 0.147 | 0.010 | 0.296 | 525 | Sialic acid biosynthesis enzyme | |
| DNAJB1 | 0.237 | 0.036 | 0.341 | 628 | 0.282 | 0.030 | 0.432 | 302 | Protein-folding co-chaperone | Protein quality control and autophagy |
| SQSTM1 | 0.071 | 0.010 | 0.141 | 1121 | 0.155 | 0.003 | 0.398 | 353 | Autophagy scaffold protein | |
| MTERF4 | 0.266 | 0.0005 | 0.876 | 117 | 0.337 | 0.021 | 0.567 | 159 | Mitochondrial ribosome factor | RNA processing, splicing and translation |
| DUS4L | 0.451 | 0.019 | 0.772 | 160 | 0.515 | 0.027 | 0.811 | 58 | tRNA modification enzyme | |
| U2SURP | 0.244 | 0.004 | 0.582 | 270 | 0.163 | 0.0005 | 0.542 | 174 | Nuclear RNA-binding splicing factor | |
| EIF2S1 | 0.222 | 0.004 | 0.523 | 321 | 0.289 | 0.003 | 0.741 | 85 | Translation initiation factor | |
| MKR N1 | 0.221 | 0.005 | 0.512 | 335 | 0.219 | 0.005 | 0.496 | 216 | Ribosome assoaciated ubiquitin ligase | |
| EIF5B | 0.263 | 0.013 | 0.496 | 361 | 0.266 | 0.004 | 0.641 | 126 | Translation initiation factor | |
| ZC3H18 | 0.292 | 0.025 | 0.465 | 403 | 0.232 | 0.038 | 0.33 | 461 | RNA stability regulatory protein | |

| | | | | | | | | | | |
|---|---|---|---|---|---|---|---|---|---|---|
| SNRPF | 0.263 | 0.018 | 0.458 | 415 | 0.31 | 0.006 | 0.684 | 113 | Spliceosomal core protein | |
| SRRM1 | 0.342 | 0.046 | 0.457 | 419 | 0.376 | 0.036 | 0.511 | 201 | Splicing regulatory protein | |
| RPAP3 | 0.234 | 0.011 | 0.455 | 422 | 0.191 | 0.024 | 0.309 | 502 | Transcriptional regulator | |
| KHDRBS1 | 0.28 | 0.034 | 0.412 | 485 | 0.283 | 0.038 | 0.401 | 344 | RNA-binding signaling adaptor | |
| PRPF38B | 0.197 | 0.017 | 0.348 | 606 | 0.226 | 0.011 | 0.447 | 278 | pre-mRNA processing factor | |
| PUS7L | 0.155 | 0.006 | 0.344 | 616 | 0.144 | 0.008 | 0.304 | 509 | mRNA pseudouridine synthase | |
| EIF2S3X | 0.202 | 0.024 | 0.327 | 655 | 0.241 | 0.038 | 0.343 | 435 | Translation initiation subunit | |
| GTF2F1 | 0.156 | 0.009 | 0.32 | 670 | 0.13 | 0.003 | 0.331 | 458 | RNA polymerase II factor | |
| EIF2A | 0.159 | 0.022 | 0.263 | 808 | 0.168 | 0.020 | 0.288 | 542 | Translation initiation factor | |
| SF3B3 | 0.174 | 0.033 | 0.258 | 831 | 0.188 | 0.032 | 0.279 | 567 | Core spliceosomal protein | |
| KARS1 | 0.155 | 0.022 | 0.257 | 837 | 0.265 | 0.021 | 0.443 | 286 | Lysyl-tRNA synthetase enzyme | |
| EIF1AX | 0.141 | 0.017 | 0.251 | 856 | 0.17 | 0.011 | 0.334 | 454 | Translation initiation factor | |
| PABPC4 | 0.159 | 0.041 | 0.22 | 931 | 0.32 | 0.034 | 0.463 | 249 | Poly(A)-binding protein | |
| RPLP2 | 0.129 | 0.037 | 0.185 | 1033 | 0.221 | 0.031 | 0.338 | 446 | Ribosomal structural protein | |
| DDX23 | 0.101 | 0.042 | 0.139 | 1125 | 0.196 | 0.034 | 0.435 | 297 | RNA helicase spliceosome factor | |
| CMTM3 | 0.657 | 0.034 | 0.964 | 90 | 1.06 | 0.011 | 2.095 | 1 | Membrane signaling protein | Signaling pathways |
| CCNQ | 0.245 | 0.024 | 0.395 | 513 | 0.241 | 0.035 | 0.351 | 426 | Transcriptional regulator | |
| PTPA | 0.19 | 0.038 | 0.27 | 780 | 0.226 | 0.031 | 0.314 | 484 | Phosphatase activator | |
| FAM50A | 0.222 | 0.002 | 0.585 | 268 | 0.104 | 0.050 | 0.136 | 858 | Nuclear transcriptional regulator | Transcription and gene expression |
| ZFP236 | 0.359 | 0.029 | 0.551 | 292 | 0.099 | 0.030 | 0.152 | 846 | Zinc-finger transcription factor | |
| SUPT5H | 0.183 | 0.011 | 0.355 | 586 | 0.151 | 0.021 | 0.251 | 627 | Transcription elongation regulator | |
| NKAPD1 | 0.219 | 0.040 | 0.304 | 701 | 0.227 | 0.032 | 0.412 | 327 | Nuclear RNA-binding regulatory factor | |
| TDRD3 | 0.158 | 0.019 | 0.27 | 782 | 0.08 | 0.029 | 0.123 | 866 | RNA-binding transcriptional co-regulator | |
| MORF4L2 | 0.157 | 0.049 | 0.206 | 969 | 0.138 | 0.038 | 0.296 | 524 | Chromatin remodeling regulator | |
| POLR3F | 0.085 | 0.032 | 0.127 | 1147 | 0.173 | 0.003 | 0.43 | 305 | RNA polymerase III subunit | |
| LRATD1 | 0.15 | 0.047 | 0.199 | 992 | 0.151 | 0.037 | 0.197 | 760 | Putative membrane protein | Miscellaneous |
