## Supplemental Table 1 for "Evolutionarily conserved temperature dependence leads to loss of protein O-GlcNAc in mammalian hypothermia"

**Supplementary Table 1**. Proteins with significantly decreased solubility ($log_2FC < 0$, $p < 0.05$) identified by PISA assay in cells cultured at 30°C vs 37°C and at 30°C vs 30°C + TMG.

| | 30°C vs 37°C | | | | 30°C vs 30°C ± TMG | | | | | |
|---|---|---|---|---|---|---|---|---|---|---|
| **Protein** | **Log₂ FC** | **p value** | **Score** | **Rank** | **Log₂ FC** | **p value** | **Score** | **Rank** | **Function** | **Functional group** |
| DNAJC15 | -0.37 | 0.002 | 1.03 | 72 | -0.27 | 0.004 | 0.65 | 122 | Mitochondrial co-chaperone | Mitochondrial function and OXPHOS |
| ATP8 | -0.49 | 0.022 | 0.81 | 141 | -0.46 | 0.033 | 0.68 | 112 | ATP synthase membrane subunit | |
| TOMM5 | -0.45 | 0.020 | 0.77 | 159 | -0.45 | 0.027 | 0.71 | 103 | Mitochondrial import adapter | |
| ATP5PO | -0.28 | 0.008 | 0.57 | 274 | -0.16 | 0.006 | 0.36 | 407 | ATP synthase structural subunit | |
| RBFA | -0.28 | 0.016 | 0.51 | 346 | -0.29 | 0.026 | 0.46 | 252 | Mitochondrial ribosome assembly factor | |
| UQCRC2 | -0.23 | 0.010 | 0.46 | 416 | -0.20 | 0.002 | 0.54 | 173 | Complex III structural subunit | |
| COX7C | -0.26 | 0.030 | 0.40 | 497 | -0.24 | 0.049 | 0.31 | 496 | Complex IV structural subunit | |
| MPC2 | -0.22 | 0.017 | 0.39 | 529 | -0.15 | 0.019 | 0.26 | 613 | Mitochondrial pyruvate transport | |
| ATP5F1C | -0.22 | 0.021 | 0.37 | 565 | -0.16 | 0.004 | 0.38 | 377 | ATP synthase rotor subunit | |
| COX20 | -0.20 | 0.039 | 0.28 | 744 | -0.24 | 0.016 | 0.43 | 306 | Complex IV assembly factor | |
| FUNDC1 | -0.18 | 0.031 | 0.27 | 783 | -0.17 | 0.021 | 0.28 | 565 | Mitophagy receptor protein | |
| SLC25A12 | -0.15 | 0.018 | 0.26 | 816 | -0.18 | 0.008 | 0.38 | 384 | Mitochondrial amino acid transporter | |
| GFER | -0.11 | 0.012 | 0.21 | 943 | -0.09 | 0.010 | 0.18 | 798 | Redox enzyme | |
| MRPL34 | -0.15 | 0.038 | 0.21 | 954 | -0.38 | 0.0005 | 1.26 | 15 | Mitochondrial ribosome component | |
| ATP5F1B | -0.12 | 0.018 | 0.21 | 963 | -0.11 | 0.027 | 0.17 | 805 | ATP synthase catalytic subunit | |
| BCS1L | -0.09 | 0.014 | 0.16 | 1086 | -0.13 | 0.030 | 0.19 | 766 | Complex III assembly factor | |
| EEF1A2 | -0.66 | 0.041 | 0.92 | 102 | -0.73 | 0.016 | 1.32 | 12 | Translation elongation factor | RNA processing, splicing and control |
| NUP58 | -0.22 | 9.59E-05 | 0.86 | 120 | -0.35 | 3.70E-05 | 1.54 | 4 | Nuclear pore transport | |
| XAB2 | -0.28 | 0.001 | 0.80 | 147 | -0.17 | 0.010 | 0.34 | 438 | Pre-mRNA splicing factor | |
| PRPF40B | -0.23 | 0.021 | 0.39 | 517 | -0.22 | 0.026 | 0.34 | 431 | mRNA splicing factor | |
| QTRT2 | -0.18 | 0.038 | 0.26 | 846 | -0.17 | 0.039 | 0.24 | 637 | tRNA queuosine modification enzyme | |
| YJU2B | -0.13 | 0.027 | 0.21 | 967 | -0.23 | 0.011 | 0.44 | 285 | Pre-mRNA splicing factor | |
| LSM1 | -0.12 | 0.041 | 0.17 | 1067 | -0.13 | 0.049 | 0.17 | 823 | RNA processing scaffold protein | |
| TARS1 | -0.08 | 0.017 | 0.14 | 1122 | -0.08 | 0.012 | 0.16 | 840 | tRNA charging enzyme | |

| | | | | | | | | | | |
|---|---|---|---|---|---|---|---|---|---|---|
| BCDIN3D | -0.10 | 0.045 | 0.13 | 1141 | -0.27 | 0.002 | 0.70 | 73 | RNA methyltransferase | |
| SAMD4B | -0.04 | 0.031 | 0.06 | 1199 | -0.07 | 0.011 | 0.13 | 860 | mRNA translation regulator | |
| GNG10 | -0.59 | 0.015 | 1.08 | 59 | -0.44 | 0.036 | 0.63 | 128 | G-protein signaling subunit | |
| ATF1 | -0.46 | 0.038 | 0.65 | 235 | -0.44 | 0.044 | 0.59 | 145 | cAMP-responsive transcription factor | |
| ZBED6 | -0.29 | 0.006 | 0.64 | 237 | -0.18 | 0.025 | 0.29 | 533 | Transcriptional repressor | |
| GTF2I | -0.36 | 0.018 | 0.63 | 242 | -0.28 | 0.043 | 0.38 | 379 | Transcriptional regulatory factor | |
| ZNF526 | -0.35 | 0.021 | 0.59 | 266 | -0.27 | 0.039 | 0.38 | 377 | Putative transcription regulator | |
| JAG2 | -0.23 | 0.005 | 0.53 | 305 | -0.63 | 0.048 | 0.83 | 49 | Notch signaling ligand | |
| MAPK6 | -0.29 | 0.017 | 0.52 | 324 | -0.50 | 0.002 | 1.36 | 9 | Serine/threonine signaling kinase | |
| EREG | -0.30 | 0.032 | 0.44 | 434 | -0.44 | 0.003 | 1.09 | 21 | EGFR ligand growth factor | |
| HCFC1 | -0.21 | 0.008 | 0.44 | 440 | -0.35 | 0.007 | 0.75 | 81 | Transcriptional co-regulator | Signaling and transcriptional regulation |
| ZFP212 | -0.28 | 0.027 | 0.43 | 452 | -0.32 | 0.012 | 0.62 | 133 | Putative transcription regulator | |
| SOCS4 | -0.25 | 0.032 | 0.38 | 544 | -0.15 | 0.026 | 0.24 | 641 | Cytokine signaling suppressor | |
| LTBR | -0.19 | 0.014 | 0.34 | 617 | -0.16 | 0.011 | 0.31 | 503 | TNF superfamily receptor | |
| MED16 | -0.16 | 0.017 | 0.29 | 734 | -0.13 | 0.036 | 0.19 | 767 | Transcriptional coactivator subunit | |
| YEATS2 | -0.20 | 0.046 | 0.26 | 812 | -0.18 | 0.049 | 0.23 | 664 | Chromatin acetylation regulator | |
| ATXN1 | -0.19 | 0.046 | 0.25 | 857 | -0.33 | 0.008 | 0.70 | 106 | Transcriptional regulation | |
| CHD6 | -0.14 | 27 | 0.21 | 950 | -0.21 | 0.013 | 0.40 | 346 | Chromatin remodeling ATPase | |
| IFNGR1 | -0.14 | 0.042 | 0.19 | 1018 | -0.15 | 0.006 | 0.32 | 476 | IFN-γ immune receptor | |
| FOXJ2 | -0.10 | 0.030 | 0.14 | 1118 | -0.24 | 0.007 | 0.52 | 193 | Forkhead transcription factor | |
| MKNK2 | -0.06 | 0.033 | 0.09 | 1177 | -0.16 | 0.040 | 0.22 | 684 | Translation-regulating kinase | |
| MISP3 | -0.74 | 0.001 | 2.09 | 10 | -0.47 | 0.007 | 1.00 | 28 | Putative spindle positioning protein | |
| DLG5 | -0.27 | 0.0002 | 0.99 | 82 | -0.11 | 0.016 | 0.20 | 761 | Cell polarity scaffold | |
| CTNNAL1 | -0.63 | 0.038 | 0.90 | 106 | -0.21 | 0.050 | 0.27 | 578 | Cytoskeleton scaffold protein | |
| MICAL2 | -0.27 | 0.0006 | 0.86 | 122 | -0.12 | 0.005 | 0.28 | 569 | Cytoskeletal dynamics | Membrane organization, trafficking and cytoskeleton |
| SLC8A1 | -0.34 | 0.020 | 0.58 | 271 | -0.46 | 0.033 | 0.68 | 114 | $Na^+/Ca^{2+}$ exchanger | |
| LRRC8E | -0.30 | 0.015 | 0.55 | 290 | -0.15 | 0.002 | 0.41 | 332 | Anion channel subunit | |
| CTTNBP2 | -0.26 | 0.009 | 0.53 | 314 | -0.37 | 0.044 | 0.50 | 211 | Actin scaffold protein | |

| | | | | | | | | | | |
|---|---|---|---|---|---|---|---|---|---|---|
| TSPAN31 | -0.22 | 0.013 | 0.41 | 493 | -0.25 | 0.024 | 0.40 | 348 | Transmembral signaling protein | |
| OCIAD1 | -0.17 | 0.012 | 0.33 | 657 | -0.17 | 0.011 | 0.34 | 445 | Membrane-bound signaling regulator | |
| DNAH12 | -0.23 | 0.044 | 0.31 | 688 | -0.33 | 0.033 | 0.49 | 219 | Microtubule motor ATPase | |
| MEGF8 | -0.20 | 0.045 | 0.27 | 778 | -0.42 | 0.015 | 0.77 | 76 | Single pass membrane protein | |
| ATP1B3 | -0.18 | 0.035 | 0.27 | 790 | -0.24 | 0.002 | 0.64 | 127 | $Na^+/K^+$ pump subunit | |
| RAB27B | -0.16 | 0.022 | 0.26 | 833 | -0.48 | 0.042 | 0.66 | 120 | Endocytic trafficking GTPase | |
| TRAPPC9 | -0.14 | 0.016 | 0.25 | 868 | -0.14 | 0.030 | 0.22 | 698 | Vesicle trafficking component | |
| SCAMP5 | -0.17 | 0.042 | 0.24 | 882 | -0.38 | 0.027 | 0.59 | 149 | Vesicle trafficking membrane protein | |
| TMEM134 | -0.11 | 0.007 | 0.24 | 892 | -0.14 | 0.023 | 0.22 | 687 | Predicted membrane protein | |
| DNAAF2 | -0.11 | 0.013 | 0.21 | 959 | -0.16 | 0.035 | 0.23 | 665 | Dynein assembly factor | |
| SNAP29 | -0.07 | 0.027 | 0.10 | 1169 | -0.14 | 0.005 | 0.31 | 482 | SNARE-mediated fusion adaptor | |
| NPRL2 | -0.30 | 0.002 | 0.82 | 137 | -0.30 | 0.007 | 0.65 | 125 | mTOR nutrient sensing | Metabolism, lipids, nutrients and small-molecule processing |
| RFESD | -0.26 | 0.001 | 0.79 | 151 | -0.19 | 0.012 | 0.37 | 399 | Putative iron–sulfur enzyme | |
| IP6K2 | -0.32 | 0.005 | 0.75 | 172 | -0.19 | 0.041 | 0.26 | 607 | Inositol phosphate kinase | |
| LPIN1 | -0.20 | 0.003 | 0.50 | 353 | -0.19 | 0.007 | 0.40 | 340 | Lipid metabolism | |
| CBR4 | -0.28 | 0.022 | 0.46 | 413 | -0.26 | 0.006 | 0.58 | 146 | Mitochondrial lipid reductase | |
| PFKFB1 | -0.30 | 0.034 | 0.44 | 441 | -0.29 | 0.004 | 0.70 | 108 | Glycolysis regulatory enzyme | |
| ALG2 | -0.24 | 0.019 | 0.42 | 480 | -0.25 | 0.030 | 0.38 | 371 | Protein glycosylation enzyme | |
| MOCS1 | -0.17 | 0.008 | 0.36 | 584 | -0.14 | 0.002 | 0.37 | 382 | Cofactor in molybdenum biosynthesis | |
| FADS1 | -0.21 | 0.026 | 0.34 | 630 | -0.36 | 0.024 | 0.59 | 144 | Fatty acid desaturase | |
| PNPLA2 | -0.24 | 0.044 | 0.33 | 652 | -0.28 | 0.013 | 0.53 | 184 | Lipid droplet lipase enzyme | |
| GCAT | -0.19 | 0.022 | 0.32 | 673 | -0.17 | 0.035 | 0.25 | 636 | Glycine metabolism enzyme | |
| CYP2J6 | -0.17 | 0.016 | 0.31 | 697 | -0.19 | 0.004 | 0.45 | 274 | Lipid oxidation enzyme | |
| ACOX1 | -0.19 | 0.027 | 0.29 | 728 | -0.12 | 0.048 | 0.16 | 841 | Peroxisomal fatty acid oxidation enzyme | |
| ADPGK | -0.16 | 0.020 | 0.28 | 756 | -0.17 | 0.022 | 0.29 | 546 | Glucose phosphorylation enzyme | |
| FDFT1 | -0.17 | 0.028 | 0.26 | 834 | -0.23 | 0.003 | 0.55 | 168 | Sterol biosynthesis enzyme | |
| DHRS7 | -0.16 | 0.031 | 0.25 | 864 | -0.17 | 0.019 | 0.29 | 543 | Redox lipid enzyme | |
| CYB561 | -0.16 | 0.037 | 0.23 | 916 | -0.16 | 0.004 | 0.38 | 380 | Transmembrane reductase | |
| PGS1 | -13 | 0.020 | 0.22 | 922 | -0.22 | 0.002 | 0.59 | 148 | Cardiolipin biosynthesis enzyme | |

| | | | | | | | | | | |
|---|---|---|---|---|---|---|---|---|---|---|
| CYB5R4 | -0.15 | 0.032 | 0.22 | 925 | -0.12 | 0.017 | 0.22 | 700 | NADH redox enzyme | |
| SLC35A3 | -0.14 | 0.037 | 0.20 | 1002 | -0.14 | 0.026 | 0.27 | 680 | Golgi nucleotide sugar transporter | |
| HMGCL | -0.09 | 0.021 | 0.16 | 1092 | -0.11 | 0.014 | 0.20 | 762 | Ketone body synthesis enzyme | |
| HYCC2 | -0.07 | 0.043 | 0.09 | 1179 | -0.11 | 0.018 | 0.18 | 788 | PI4KA lipid kinase complex subunit | |
| RTEL1 | -0.30 | 0,0004 | 1.00 | 79 | -0.57 | 0.025 | 0.91 | 36 | DNA helicase stability | Genome stability, DNA repair and cell cycle |
| MBD4 | -0.41 | 0.010 | 0.82 | 139 | -0.41 | 0.011 | 0.81 | 57 | DNA mismatch repair enzyme | |
| REV1 | -0.35 | 0.009 | 0.72 | 191 | -0.33 | 0.045 | 0.44 | 291 | DNA damage tolerance polymerase | |
| LIN37 | -0.14 | 0.010 | 0.28 | 759 | -0.19 | 0.007 | 0.42 | 317 | Cell cycle repressor | |
| RTEL1 | -0.30 | 0,0004 | 1.00 | 79 | -0.57 | 0.025 | 0.91 | 36 | DNA helicase stability | Miscellaneous |
| BCL2L1 | -0.23 | 0.013 | 0.42 | 470 | -0.12 | 0.049 | 0.16 | 844 | Anti-apoptotic regulator | |
| ANKRD10 | -0.24 | 0.019 | 0.42 | 473 | -0.28 | 0.028 | 0.44 | 293 | Putative scaffold protein | |
| CACFD1 | -0.20 | 0.010 | 0.40 | 495 | -0.16 | 0.027 | 0.25 | 623 | Predicted calcium handling protein | |
| PILRA | -0.18 | 0.020 | 0.31 | 686 | -0.14 | 0.022 | 0.24 | 658 | Inhibitory immune receptor | |
| MPND | -0.16 | 0.019 | 0.27 | 798 | -0.23 | 0.022 | 0.38 | 370 | MPN domain-containing protein | |
| ABCC4 | -0.15 | 0.030 | 0.22 | 919 | -0.12 | 0.037 | 0.17 | 829 | ATP-dependent exporter | |
| SELENOS | -0.15 | 0.047 | 0.20 | 1007 | -0.24 | 0.011 | 0.47 | 241 | ER protein quality control | |
| APMAP | -0.14 | 0.042 | 0.19 | 1024 | -0.18 | 0.003 | 0.45 | 275 | Adipocyte membrane protein | |
