## Supplementary figures and images for "Evolutionarily conserved temperature dependence leads to loss of protein O-GlcNAc in mammalian hypothermia"

### Supplemental figures

#### Supplementary Figure 1

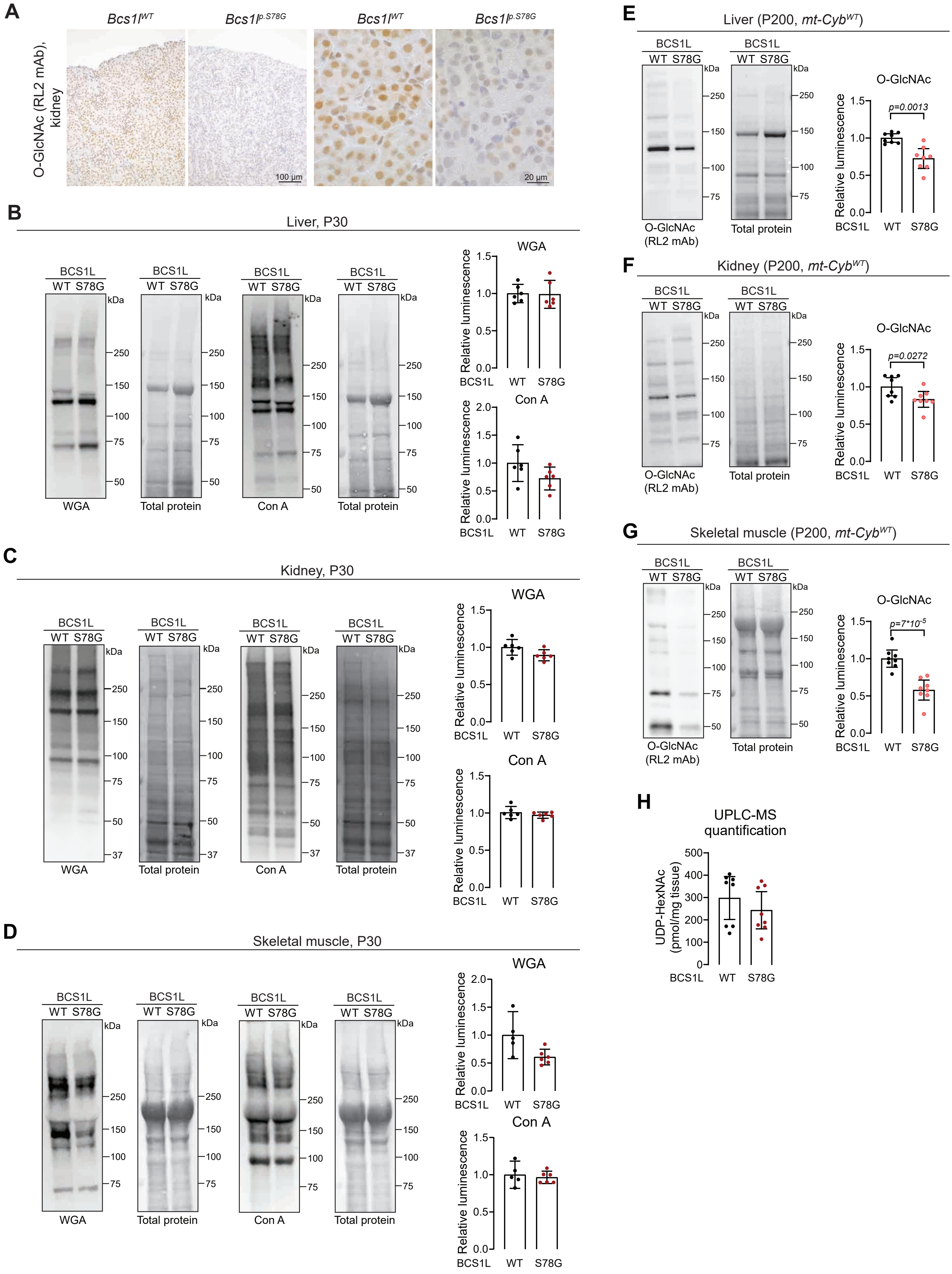

#### Supplementary Figure 2

A

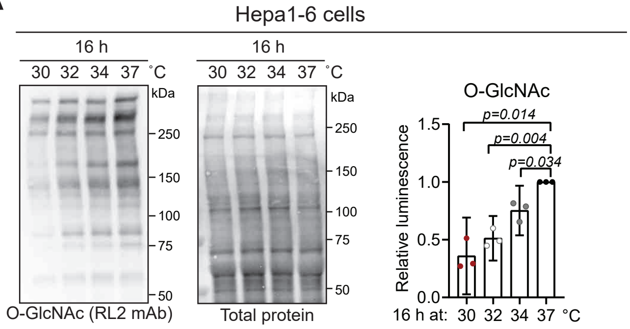


B

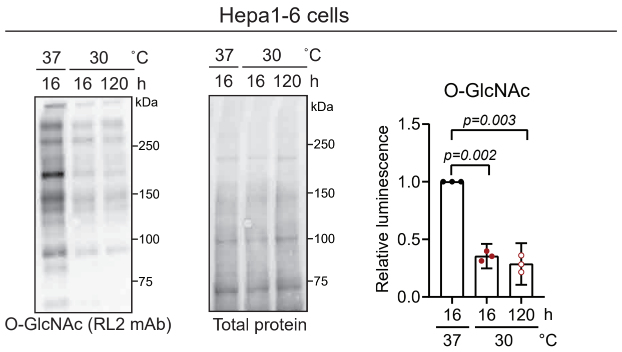


C

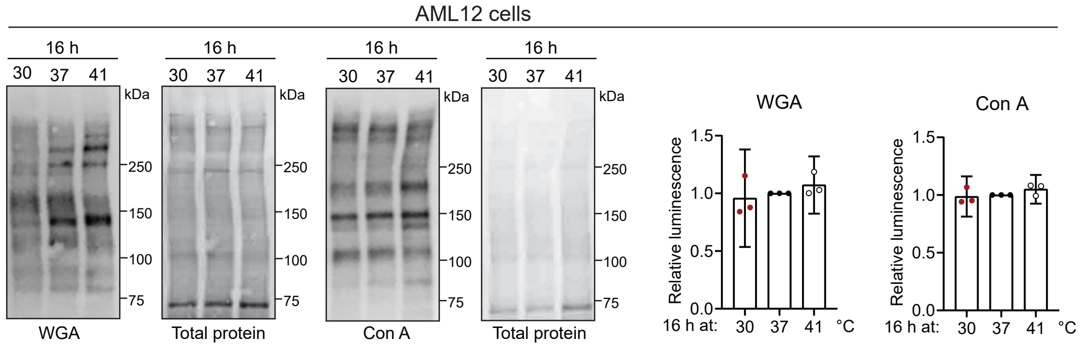

#### Supplementary Figure 3

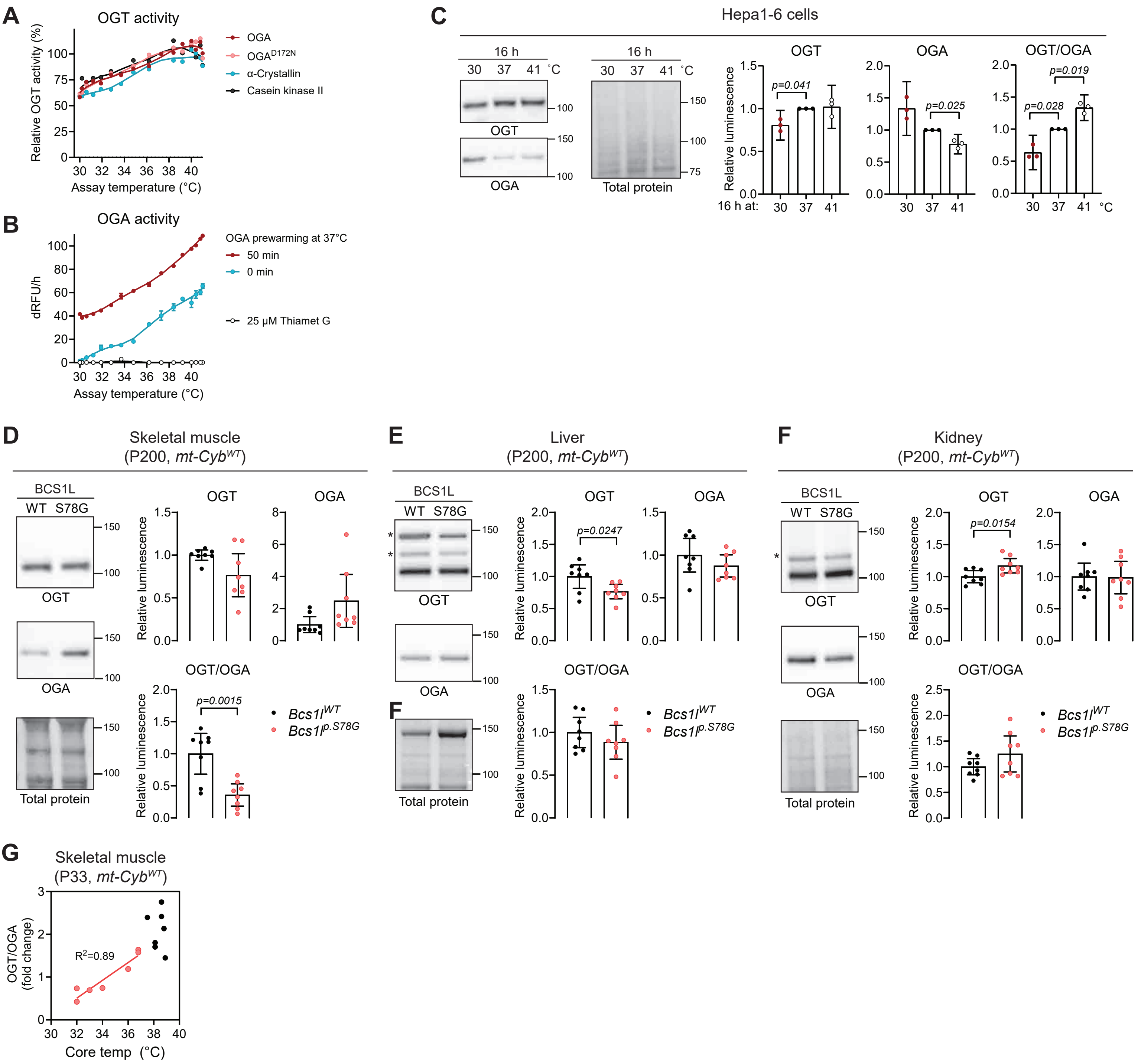

#### Supplementary Figure 4

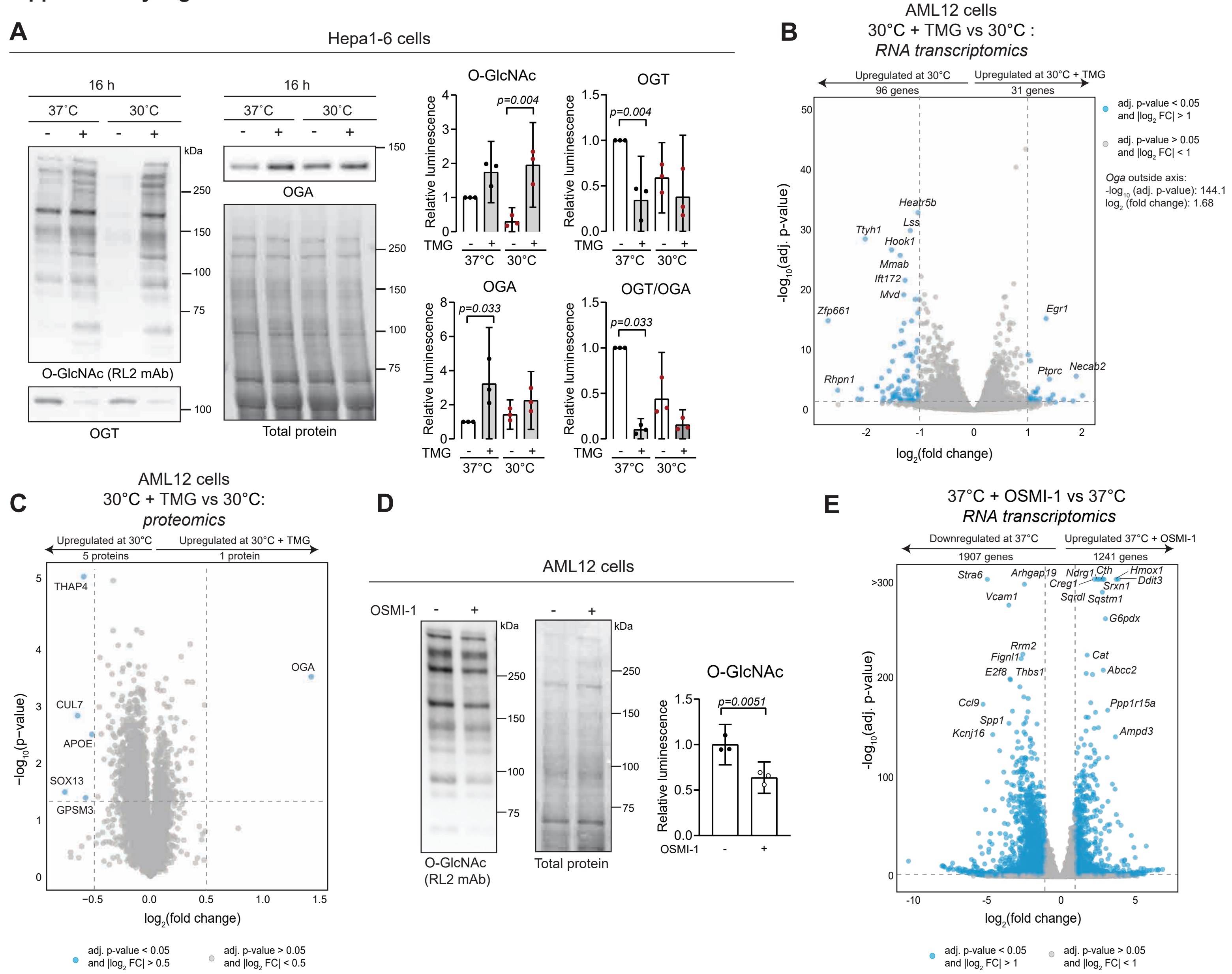
